# Genome scans of dog behavior implicate a gene network underlying psychopathology in mammals, including humans

**DOI:** 10.1101/2020.07.19.211078

**Authors:** Isain Zapata, Erin E. Hecht, James A. Serpell, Carlos E. Alvarez

## Abstract

Genetic studies show a general factor associated with all human psychopathology and strongly correlated with personality and intelligence, but its basis is unknown. We performed genome scans of 17 normal and problem behaviors in three multi-breed dog cohorts. 21 of 90 mapped loci were supported for the same, or a related, trait in a second cohort. Several of those loci were also associated with brain structure differences across breeds; and six of the respective top-candidate genes are also associated with human brain structure and function. More broadly, the geneset of canine behavioral scans is supported by enrichment for genes mapped for human behavior, personality, cognition, psychopathology and brain structure. The biology implicated includes, neurogenesis, axon guidance, angiogenesis, brain structure, alternative splicing, disease association, Hox-family transcription factors, and subiculum expression. Because body size and behavior are correlated in dogs, we isolated the effect of body size in the dog mapping and in the comparative human UK Biobank analyses. Our dog findings are consistent with pleiotropy of diverse brain traits with energy metabolism and growth, and suggest behavioral variations often affect neurogenesis. There is support for such pleiotropy in humans and well-powered genetic studies of human psychiatric traits consistently implicate neurogenesis. We propose a genetic network which underlies neuron birth and development throughout life is associated with evolutionary adaptation of behavior and the general psychopathology factor. This understanding has implications for genetic and environmental contributions to psychiatric disease. We discuss how canine translational models can further accelerate the study of psychopathology.

**Author summary:** We genetically mapped diverse normal and problem behaviors in dogs. The well-established approach we used is ideally suited for finding variation that is common across dog breeds and for pin-pointing the most likely gene candidates. Our analysis of the genes implicated at 90 genome regions shows they are enriched for i) genes mapped for diverse brain functions and pathologies in humans; ii) genes involved in brain development throughout life; and iii) footprints of evolution in dogs, humans and other animals. We propose that is consistent with evolutionary conservation of the general genetic factor of mental health in humans, which is correlated with personality and intelligence. The implications are that this super-network of genes is preferentially targeted by evolutionary adaptation for behavior and that its dysregulation increases risk of mental health disorders.

## Introduction

Beginning in 1916 and peaking in the last decade, twin and family studies have revealed a general genetic factor “p” that underlies risk for all human psychopathology^1-3^; and which is strongly correlated with personality and intelligence^4,5^. This understanding is consistent with more recent genome wide association studies (GWASs) in humans^2^ and dogs^6,7^. Human cross-disorder psychiatric GWASs showed high levels of polygenicity and genetic-relatedness^8^. 109/146 of those mapped loci (∼75%) were pleiotropic for psychiatric traits and were enriched for neurodevelopmental genes^8^. Those findings are consistent with cross-trait studies in dogs^6^, and with human GWASs of individual traits involving diverse brain functions and structure. However, essentially all variants mapped for human complex traits have minute or small effect sizes. Thus, much of the current focus is on polygenic risk scores, and few individual risk variations have clinical or experimental utility. Biological understanding is more likely to be derived from rare large-effect variations. In contrast to human, canine behavioral pleiotropy manifests in many moderate-to-large effect variations that are common across breeds^6,7^. These present powerful and unique opportunities for biological dissection and medical translation^9,10^.

## Results

### Interbreed genome scanning of behavior, body size and lifespan

Interbreed scanning has two great advantages: i) it allows mapping of variations that are commonly fixed in different breeds, and ii) it has the effect of fine mapping due to the breaking down of linkage disequilibrium (LD) on both sides of functional variations^7,11^. As we and others have done before^6,7^, we performed GWASs using breed averages of 17 C-BARQ behavioral phenotypes (Suppl. Table S1; ref. ^12^). Fourteen of these C-BARQ traits had average interbreed-heritabilities of 0.51 ±0.12 (SD; *h*^2^ range =0.27 for non-social fear to 0.77 for trainability)^6^. We also mapped body mass and lifespan (Suppl. Text). We used genotypes from 3,752 dogs total in three cohorts of partially-overlapping breed make-up to conduct separate GWASs of each trait in each cohort (29 breeds, *n*=444; 37 breeds, *n*=423; and 50 breeds, *n*=2885)^13-15^. Cross-cohort analysis provides quasi-replication and reduces false positives due to population structure and latent variables. We measured single-marker association for each trait, correcting for population structure within each cohort by using linear mixed models in GEMMA^16^. The inflation factor (λ) had a narrow range with an average of 1.14, slightly above the 1.05-1.1 considered benign (Suppl. Text). In total, we found genome wide significant associations at 90 loci across all behavioral traits (Fig. 1A; Table 1; Suppl. Data 1, Tables S2-6)^13-15^. Eleven loci were supported for the same or a related trait in at least a second cohort – here or in our previous mapping^7^ – or for different behaviors. Ten additional loci were supported by another interbreed GWAS^6^. That quasi-replication mitigates possible type I error (see Discussion for support from studies with individual-level phenotypes and genotypes^17,18^). The geneset analyses below suggest it is unlikely our mapping has a high false positive rate.

**Table 1.**
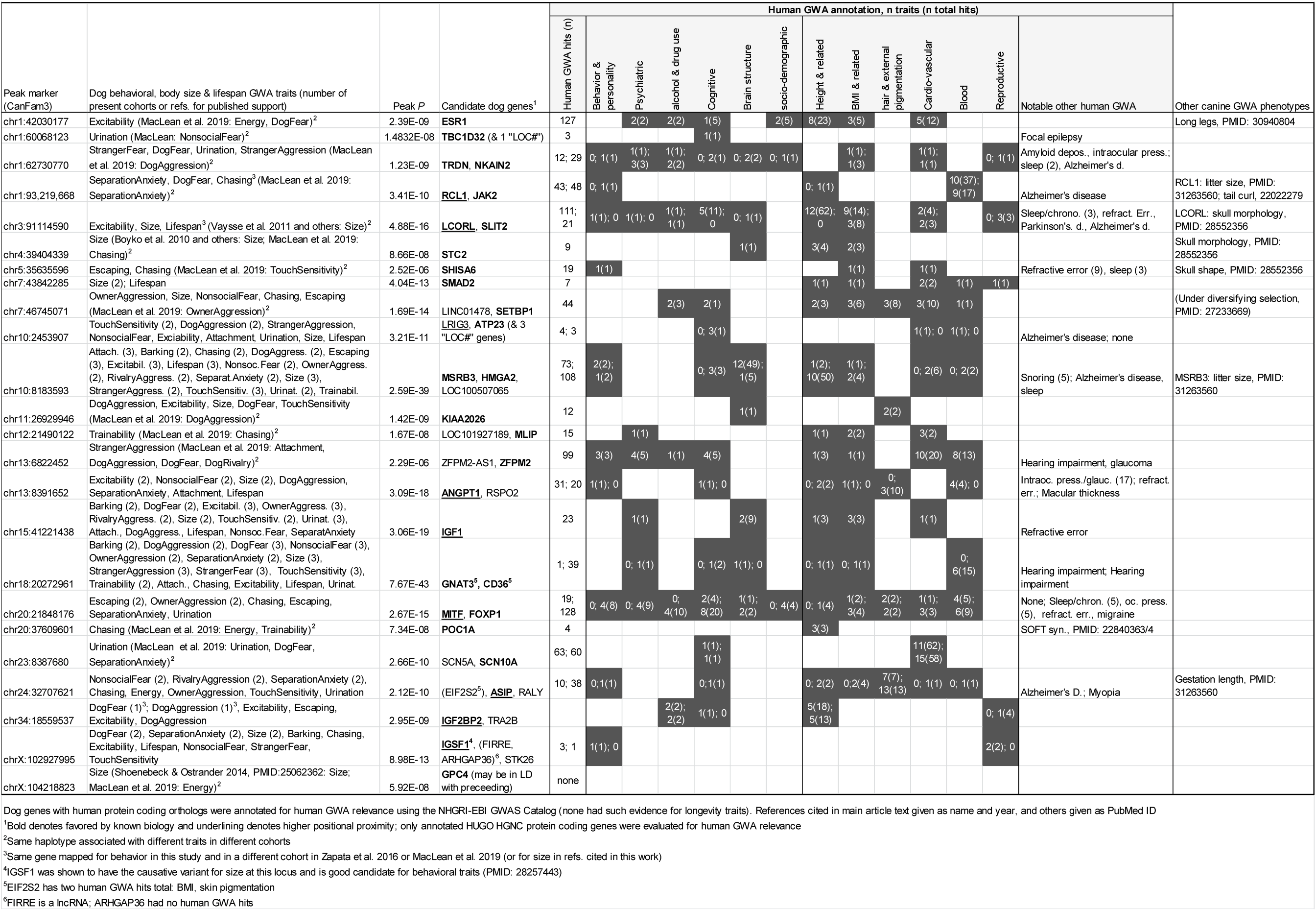
Canine interbreed GWA top confidence findings with human GWA relevance implicated in this work.

**Figure 1.**
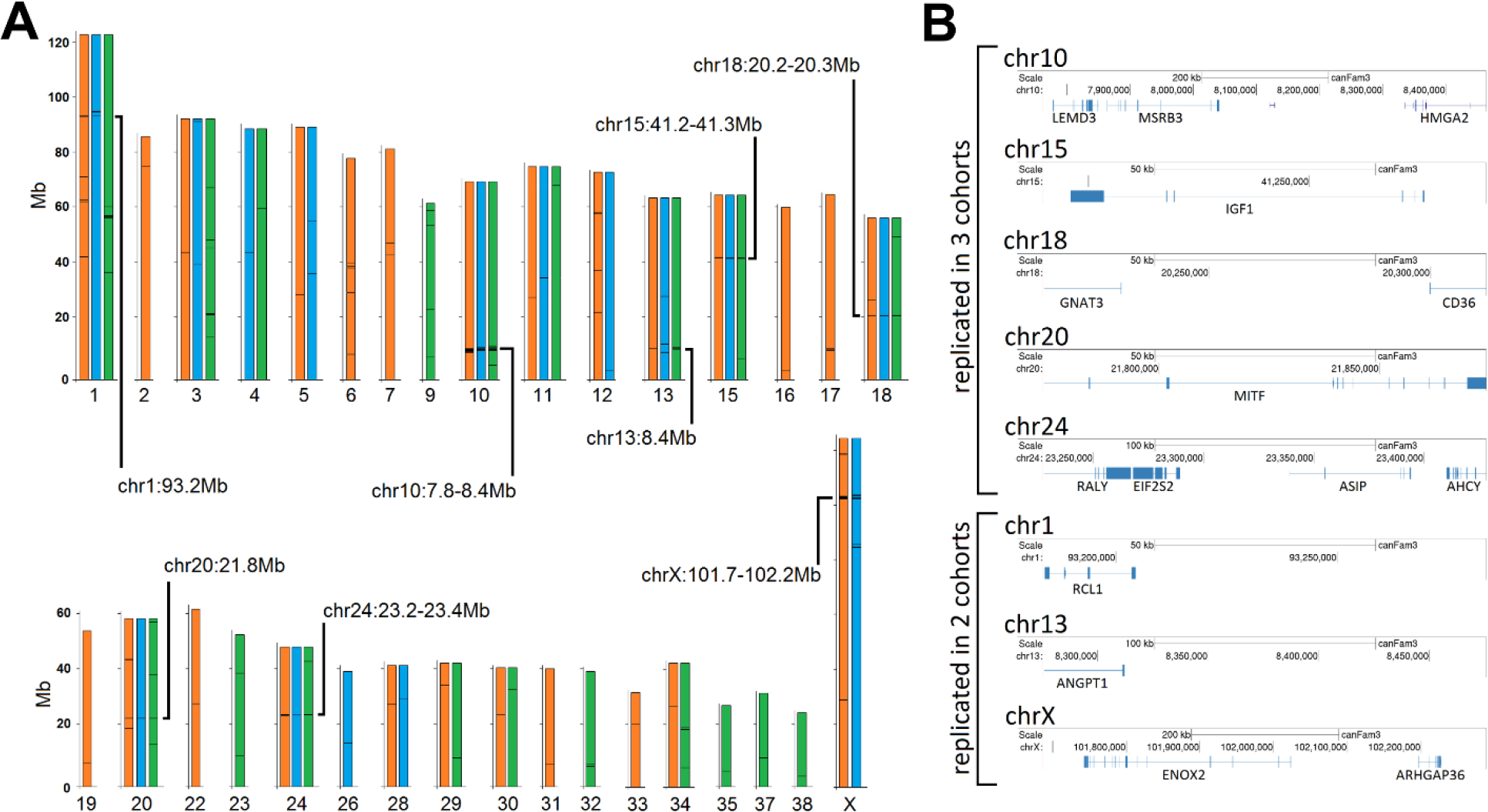
Canine behavioral mapping. (**A**) Summary of behavioral quantitative trait loci quasi-replicated in independent cohorts in this work. All loci were significant after Bonferroni adjustment. Only positive chromosomes are shown for each cohort: Vaysse et al. in orange, Boyko et al. in blue and Hayward et al. in green^13-15^. Genome coordinates are from CanFam3.1. (**B**) Mapped intervals and gene annotation for select quasi-replicated loci.

### Effect direction and amount of variation explained

To determine effect direction, we built regression models using stepwise selection of loci with significant contributions to the model (Fig. 2A). Eigen decomposition was first used to cluster the markers correlated by linkage into one. Several loci showed the presence of multiple regions with different effects on the variable. This has been reported for morphological and behavioral traits mapped to some of these loci (e.g., chr10 *MSRB3-HGMA2* locus^7,14^). The loci at chr10, chr15, chr18, chr20 and chrX were among the strongest findings. The directions of the effects are corroborated across the three cohorts (note Boyko appears opposite due to designation of minor allele; e.g., chr15/*IGF1* and body size). Many associations were significant after Bonferroni adjustment.

**Figure 2.**
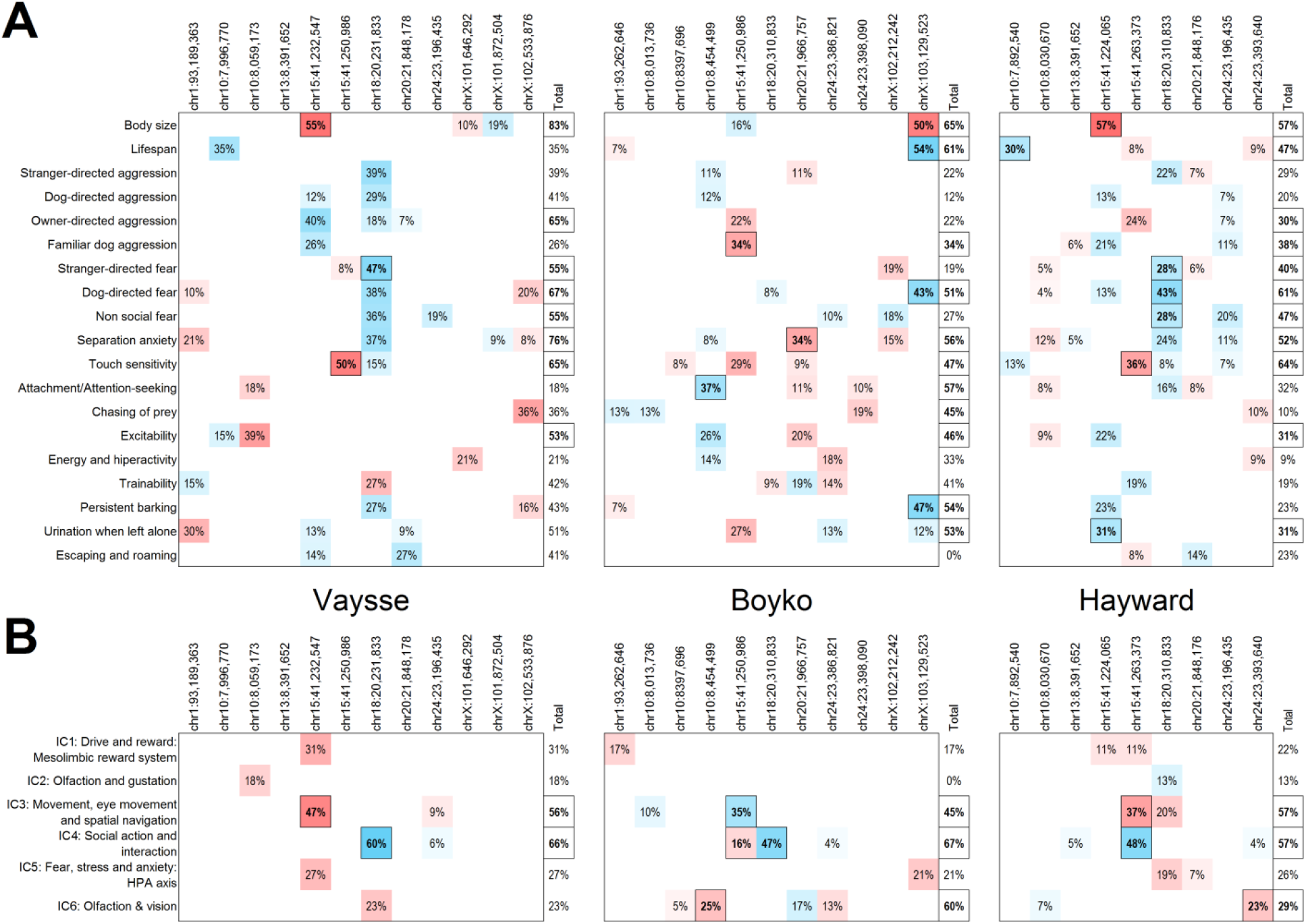
Effect directions and variance contributions. (**A**) Effect directions and variance contributions of relevant loci to behavioral traits estimated by linear regression. (**B**) Effect directions and variance contributions of relevant loci to Independent Component brain networks estimated by linear regression. Each relevant locus represents a compiled group of individual SNPs that are correlated grouped through Eigen decomposition of markers correlated by linkage. Cohorts correspond to Vaysse et al., Boyko et al. and Hayward et al., respectively^13-15^. Only significant associations are shown. Percentages correspond to *R*^*2*^ values while total indicate the overall model *R*^*2*^. Positive associations are indicated by a blue gradient while negative associations are indicated by a red gradient. Boxed and bolded values are Bonferroni significant for the cohort.

### Genetic associations with brain structure

We performed joint analyses of behavioral loci and published brain imaging data from 62 dogs of both sexes from 33 breeds^19^. Magnetic resonance imaging (MRI) data was normalized for brain volume, and used to measure gray matter differences across breeds and sexes. We tested for association using allele frequencies at our quasi-replicated loci and the factor loading coefficients of each of the six reported independent component (IC) brain networks (Fig. 2B; Suppl. Text). Note IC2, IC3 and IC5 are strongly-significantly associated, while IC1 is marginally associated with total brain volume^19^. We evaluated each association all together for all IC’s and all relevant loci in our analysis. Many associations were significant after Bonferroni adjustment. Of those, both IC2 and IC3 were associated with the chr15 and chr18 loci in all three cohorts, these associations reached individual Bonferroni significance. Multiple linear regression estimated all priority loci together account for 56%, 45% and 57% of the variance for IC2 in the three cohorts while they accounted for 66%, 67% and 57% for IC3 in the three cohorts. For IC6, two cohorts showed the variance explained was 60% and 29%. Supporting our findings, six top candidate genes at three of these loci have also been reported to be associated with differences in human brain structure: *MSRB3* and *HMGA2*, chr10 locus; *FOXP1* and *MITF*, chr20; *IGF1*, chr15; and *GNAT3*, chr18 (Table 1)^20-22^. Human alleles of those first four genes were associated with both brain structure and cognitive traits in the same study^21^.

### Trait associations, gene annotation and brain relevance of quasi-replicated loci

The four loci mapped for fear and aggression traits in multiple cohorts in our prior study^7^ (chr10:8.1, chr15, chr18 and chrX:102.1) were confirmed here and additional trait associations were identified, including brain structure and body size for all four. As we previously reported, the gene annotation of those loci shows strong behavioral relevance (Suppl. Text)^7^. The new loci on chromosomes 20 and 24 have strong behavioral candidates (Suppl. Text). Both loci are under strong evolutionary selection, but the coat-pattern alleles presumed to be under selection cannot be discerned from the present SNP genotype data^23^. Two cohorts had chr20 associations with escaping and owner-directed aggression; and single cohorts had associations with chasing, separation anxiety and separation-related urination. The chr20 locus contains three transcription factor genes: *MITF, MDFIC2* and *FOXP1*. The chr24 locus was mapped in two cohorts for nonsocial fear, rivalry aggression and separation anxiety. The genes nearest the peak marker are *RALY, EIF2S2, ASIP* and *AHCY*. Both *MITF* and agouti/*ASIP* are well known for key roles in external pigmentation, but also have strong evidence of behavioral roles. Although *MITF* is at the same locus as the prominent behavioral gene *FOXP1*, math ability, educational attainment and brain structure traits have been specifically mapped to *MITF* in human GWASs^21,24,25^. Strikingly, the targeted null-mutation of *Asip* in a wild-derived mouse strain strongly increased tameness and reduced aggression in comparison to the wild type strain^26^. Domesticated strains of several mammalian species are deficient in *Asip* and are tamer than wild type (Suppl. Text)^27-30^. The most famous example is Belyaev’s “Russian farm-foxes” which were selected for tameness for 50 generations beginning in 1959. Notably, the initial “silver foxes” tamed were silver- or black-colored Canadian red foxes later found to be null for *Asip*^31^. We found that *ASIP* mRNA is expressed in specific brain regions in humans and cattle, most prominently in the subiculum, subregions of the hippocampus, amygdala and hypothalamus, and cerebellum (Suppl. Text). These and the cited findings suggest *Asip* is one of the most important genes in domestication and the seminal tameness-variant in Belyaev’s silver foxes. This presents a major cautionary note given the vast majority of mammalian neuroscience research is conducted in C57BL/6J mice, which are null for *Asip* (“*a*” in mouse).

The following two quasi-replicated loci are among those supported by biological relevance (Suppl. Fig. S1; Suppl. Text). The haplotype on chr1, which implicates *RCL1*, was previously known to be associated with morphological and reproductive traits^14,15^. Here it was associated with chasing in two cohorts and anxiety traits in single cohorts. Human linkage and association studies showed a missense mutation in *RCL1* is associated with depression^32^. *ANGPT1* is very near the chr13 peak associated with excitability, nonsocial fear and body size in two cohorts and with dog aggression, separation anxiety, attachment and lifespan in single cohorts. This chr13 region has strong signals of evolutionary selection and interbreed differentiation in many breeds^14^, but the basis for that is unknown. A SNP within human *ANGPT1* is associated with subjective well-being in Europeans^33^.

### Biological pathway and transcription factor relevance

We identified 127 candidate genes in the 90 loci mapped for any trait. For all geneset analyses, we only used genes with a human protein-coding ortholog (HUGO HGNC^34^). To evaluate the effect of body size, we compared our findings with a similar study that controlled for body weight. MacLean and colleagues mapped breed averages of 14 C-BARQ behaviors included in this work, but designed the association analysis to remove the effect of body weight as a proxy for body size^6^. They used two genotype cohorts, one in common with this work^13^. Our single-marker geneset of 108 and their gene-based mapping geneset of 715 share 19 genes, which makes them significantly overlapping (hypergeometric *P*=6.91×10^−6^). First, we tested for enrichment in diverse curated categories using Enrichr and DAVID^35,36^. Table 2 is a summary of the pathway and tissue analyses (Suppl. Tables S7-33). Many associations are shared by the two canine behavioral GWA genesets, including the top ranking of the biological pathway axon guidance and the annotation for tissue enrichment of brain. Both were significant for various aspects of embryonic development and morphogenesis. Both were significant and similarly ranked for alternative splicing and disease mutation. Based on genome wide *in situ* hybridization data for mouse brain, the two genesets strongly overlapped for the subiculum and its subregions.

**Table 2.** Pathway & tissue enrichment of canine behavioral GWA.

|  | This study |  | Body-mass corrected<sup>3</sup> |  |
| --- | --- | --- | --- | --- |
|  | Rank | Bonferroni *P* | Rank | Bonferroni *P* |
| Biol. Pathways (KEGG, hum)<sup>1</sup> |  |  |  |  |
| <u>Axon guidance</u> | 1 | **1.76E-02** | 1 | **6.69E-04** |
| Biol. Pathways (gene ontology, GO)<sup>2</sup> |  |  |  |  |
| Embryo development | 1 | **4.32E-03** | 184 | 1.00E+00 |
| <u>Any term re:morphogenesis</u> | 1 | **4.32E-03** | 1 | **1.02E-06** |
| Any term containing "epithelial" | 3 | 7.84E-02 | 37 | 2.80E-01 |
| Any term containing "proliferation" | 3 | 7.84E-02 | 202 | 1.00E+00 |
| Any term containing "growth" | 10 | 7.47E-01 | 50 | **2.62E-02** |
| Any term re:neuronal development | 54 | 1.00E+00 | 4 | **2.40E-05** |
| Annotation: keyword up (Swiss-Prot)<sup>2</sup> |  |  |  |  |
| <u>Alternative splicing</u> | 1 | **4.23E-05** | 1 | **9.79E-36** |
| <u>Disease mutation</u> | 2 | **4.05E-02** | 5 | **4.10E-04** |
| Annotation: tissue up (Uniprot)<sup>2</sup> |  |  |  |  |
| Brain | 1 | 9.97E-01 | 1 | **1.12E-11** |
| Tissue expression (hum. BioGPS enriched)<sup>1</sup> |  |  |  |  |
| Uterus | 1 | **1.22E-04** | 10 | 2.03E-01 |
| Prefrontal Cortex | 7 | 4.81E-01 | 1 | **1.80E-04** |
| Fetal Brain | 54 | 1.00E+00 | 2 | **4.96E-03** |
| Brain region up (mouse Allen Brain Atlas)<sup>1</sup> |  |  |  |  |
| <u>molecular layer of Subiculum</u> | 1 | **1.19E-02** | 17 | **4.53E-03** |
| <u>Subiculum</u> | 2 | **2.69E-02** | 7 | **8.81E-04** |
| nucleus of the stria terminalis (Mv) | 3 | **2.73E-02** | 858 | 6.77E-01 |
| <u>superficial stratum of Subiculum</u> | 4 | **2.80E-02** | 25 | **1.33E-02** |
| parastrial preoptic nucleus | 5 | **2.96E-02** | 1233 | 8.63E-01 |
| mantle zone of THyB-D | 6 | **3.15E-02** | 1479 | 9.22E-01 |
| <u>mantle zone of Subiculum</u> | 7 | **3.36E-02** | 27 | **1.40E-02** |
| intermediate stratum of the VAP | 8 | **3.60E-02** | 734 | 5.80E-01 |
| dorsal part of THyB | 9 | **3.87E-02** | 1678 | 9.52E-01 |
| Ventromedial hypothalamic nucleus | 10 | **4.04E-02** | 1209 | 8.54E-01 |
| layer 5 of PaS | 11 | **4.09E-02** | 84 | **6.57E-02** |
| Ventromedial hypothalamic nuc. (C) | 12 | **4.19E-02** | 613 | 4.75E-01 |
| Uvula (IX) | 13 | **4.58E-02** | 1968 | 9.87E-01 |
| Subiculum, dorsal part | 33 | 9.10E-02 | 1 | **9.05E-05** |
| Subiculum, dorsal part, molecular layer | 111 | 1.72E-01 | 2 | **4.88E-04** |
| Subiculum, dorsal part, stratum radiatum | 100 | 1.67E-01 | 3 | **5.06E-04** |
| Striatum dorsal region | 1435 | 8.77E-01 | 4 | **6.07E-04** |
| Caudoputamen | 1752 | 9.73E-01 | 5 | **7.59E-04** |
| Field CA1 | 255 | 2.67E-01 | 6 | **7.81E-04** |
| striatum (corpus striatum) | 1055 | 7.00E-01 | 8 | **9.09E-04** |
| mantle zone of Striatum | 1283 | 7.82E-01 | 9 | **1.01E-03** |
| Anterior cingulate area, ventral part, 6b | 475 | 3.74E-01 | 10 | **1.01E-03** |
| TF-KO gene expression changes<sup>1</sup> |  |  |  |  |
| <u>LMX1B</u> | 1 | **1.64E-03** | 19 | **1.94E-02** |
| <u>MECP2</u> | 2 | **2.72E-03** | 1 | **1.35E-04** |

<sup>1</sup>Enrichr analysis, <sup>2</sup>DAVID analysis, <sup>3</sup>MacLean et al. 2019
Terms significant in both studies are italicized and underlined; $P_{Bonf} < 0.05$ are in bold

Approximately 100 transcription factor binding sites were significantly enriched for our geneset (Suppl. Table S25). The top 20 binding sites correspond to 26 transcription factors enriched for developmental processes, including 11 associated with both generation and differentiation of neurons (Table 3). Analysis of our geneset combined with MacLean et al.’s yielded 132 factors predicted by two algorithms to bind at 98% of mapped genes (Suppl. Table S34). With their overlap underlined, the top factors according to significance, beginning with the first, were <u>POU6F1</u>, <u>LHX3</u>, <u>FOXA2</u>, CDC5L, ALX1, PRX2 and MEF2A; and, according to fold-enrichment, IRF1, FOXO3, <u>LHX3</u>, FOXD3, FOXF2, <u>POU6F1</u>, <u>FOXA2</u>. Those are enriched for two classes of developmental transcription factor families: Fox and Hox (n=4, *P*_*Adj*_=1.11×10^−5^, and n=3, *P*_*Adj*_=0.153, respectively; Enrichr).

**Table 3.**
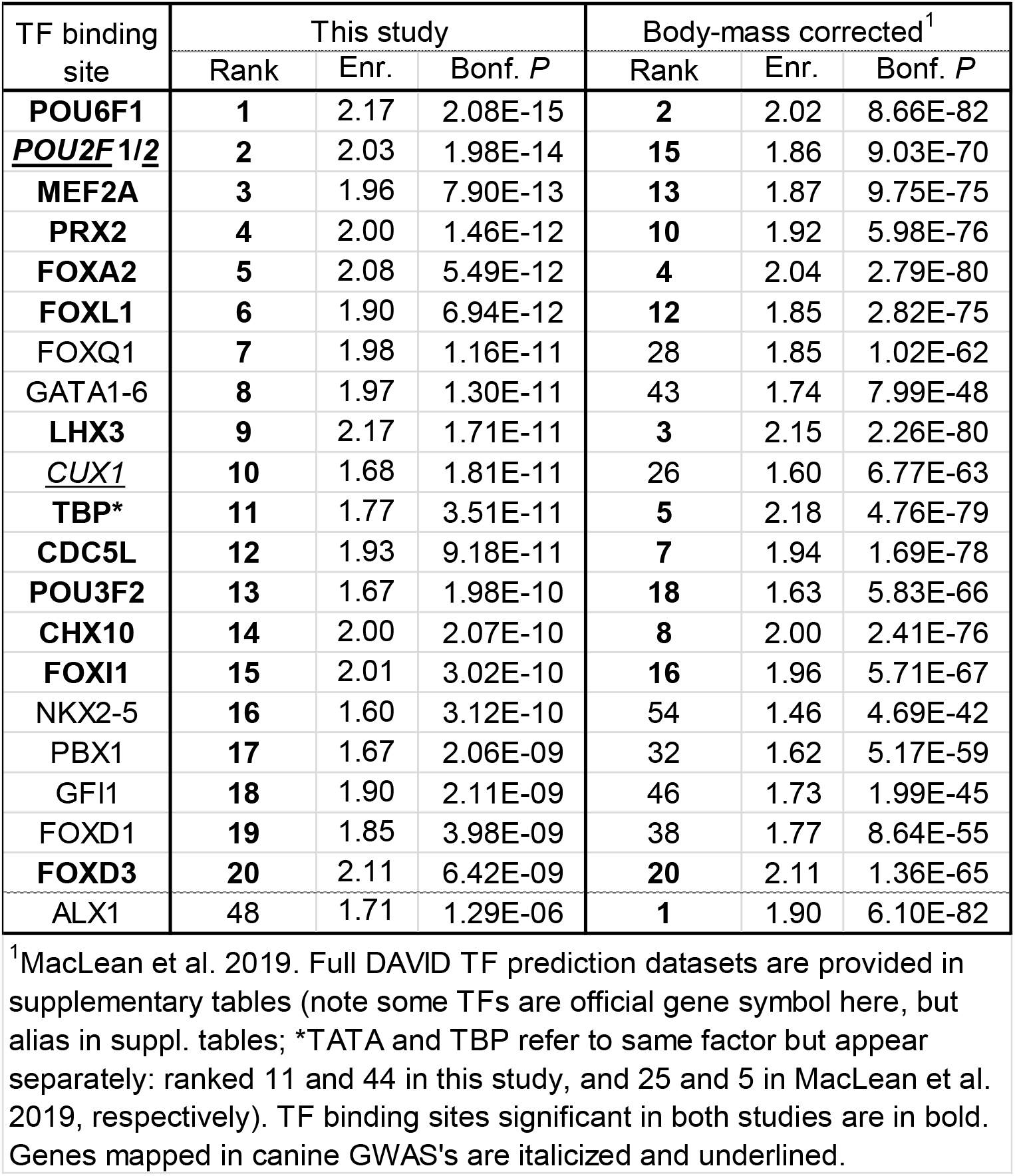
Transcription factor binding site prediction.

### Human GWA relevance

Table 4 summarizes results of the geneset analyses of human genetics. We first tested the compiled set of all GWASs with *n*>50,000 (ref. ^37^). The MacLean geneset was enriched for the categories Activities, Environment, Cognition and Reproduction (Psychiatric was suggestive), of which Activities and Environment remained significant after Bonferroni correction (Suppl. Table S35). The small Social Interactions set (379 genes) was omitted from the main analysis, but was suggestive (*P*=7.95×10^−3^). Our far lower-powered geneset was not significant for any, but had the smallest *P* value for the same category as MacLean’s and combining the two improved the significance of all positive categories. We tested the two dog genesets in parallel against three datasets of human genetics (Suppl. Tables S27-33). Enrichment testing of the GWAS Catalog (NHGRI-EBI) strongly reflected the difference in our study design vs. MacLean et al.’s, who sought to remove the effect of body size (Table 4). The traits that were significant and top-ranked in both genesets were systolic blood pressure, QRS duration and pulse pressure. Analysis of the UK Biobank human GWAS dataset revealed 79 significant traits for our geneset and 181 for MacLean et al.’s. Both the UK Biobank and dbGAP analyses showed our geneset ranked higher than MacLean’s for height-related traits and vice versa for traits related to metabolism and body mass index (BMI). These patterns suggest breed-average weights are good proxies for body size in dogs, and show much of that variation was removed in MacLean et al.’s GWASs (whereas BMI ranked #1). However, the MacLean geneset contained four of our nine genes/loci quasi-replicated for behavior and associated with body size (*MSRB3, ANGPT1, IGF2BP2, IGSF1*), and had strong signal for height in the UK Biobank GWAS and dbGAP (Bonferroni *P*=1.08×10^−5^ and *P*=5.63×10^−8^, respectively).

**Table 4.** Human genetics relevance of canine behavioral GWA.

| Sources and representative traits | This study |  | Body-mass corrected<sup>1</sup> |  |
| --- | --- | --- | --- | --- |
|  | Rank | Bonf. *P* | Rank | Bonf. *P* |
| Human GWA catalog |  |  |  |  |
| Hip circumference adjusted for BMI* | 1 | **1.89E-03** | 146 | 3.01E-01 |
| Height* | 2 | **4.64E-03** | 108* | 2.49E-01 |
| <u>Systolic blood pressure</u> | 3 | **6.96E-03** | 2 | **5.45E-07** |
| <u>Menarche (age at onset)</u> | 4 | **8.61E-03** | 18 | <u>6.13E-02</u> |
| PR interval | 5 | **1.08E-02** | 169 | 3.23E-01 |
| Birth weight* | 6 | **1.27E-02** | 105* | 2.44E-01 |
| Infant length* | 7 | **1.65E-02** | 804* | 9.20E-01 |
| Heart rate response to recovery post exercise | 8 | **3.03E-02** | 375 | 7.03E-01 |
| <u>QRS duration</u> | 9 | **4.28E-02** | 13 | **2.55E-02** |
| <u>Pulse pressure</u> | 10 | **4.31E-02** | 5 | **3.19E-03** |
| Human GWA, UK Biobank |  |  |  |  |
| <u>Distance between home and job workplace 796 raw</u> | 1 | **1.45E-15** | 2 | **3.14E-50** |
| <u>Pulse wave Arterial Stiffness index 21021 raw</u> | 2 | **4.75E-09** | 3 | **2.93E-47** |
| <u>High light scatter reticulocyte percentage 30290 raw</u> | 3 | **3.73E-08** | 1 | **1.72E-57** |
| <u>Forced vital capacity 20151 raw</u> | 4 | **2.76E-07** | 75 | **1.02E-04** |
| <u>Standing height 50 raw</u> | 6 | **1.38E-06** | 60 | **1.08E-05** |
| <u>Forced expiratory volume in 1-second (best measure) 20150...</u> | 10 | **2.75E-06** | 83 | **1.96E-04** |
| <u>Whole body fat-free mass 23101 raw</u> | 15 | **1.27E-05** | 16 | **7.34E-14** |
| <u>Monocyte count 30130 raw</u> | 17 | **2.55E-05** | 7 | **4.07E-23** |
| <u>Weight 23098 raw</u> | 19 | **3.30E-05** | 41 | **1.36E-07** |
| <u>Longest period of unenthusiasm/disinterest 5375 raw</u> | 23 | **3.65E-05** | 10 | **9.56E-19** |
| <u>Freq...needing morning drink of alcohol after heavy drinking...</u> | 26 | **6.29E-05** | 6 | **1.46E-25** |
| <u>Basal metabolic rate 23105 raw</u> | 29 | **6.72E-05** | 18 | **8.50E-13** |
| <u>Inverse distance to the nearest major road 24012 raw</u> | 34 | **1.58E-04** | 11 | **1.76E-18** |
| <u>Hip circumference 49 raw</u> | 35 | **1.61E-04** | 54 | **4.55E-06** |
| <u>Birth weight 20022 raw</u> | 42 | **5.25E-04** | 163 | **2.80E-02** |
| <u>Lymphocyte count 30120 raw</u> | 45 | **5.72E-04** | 4 | **3.83E-41** |
| <u>Duration to first press of snap-button in each round 404 raw</u> | 51 | **1.19E-03** | 12 | **1.88E-18** |
| <u>Longest period of depression 4609 raw</u> | 52 | **1.48E-03** | 29 | **3.82E-09** |
| <u>Peak expiratory flow 3064 raw</u> | 54 | **1.92E-03** | 119 | **2.91E-03** |
| <u>Red blood cell count 30010 raw</u> | 58 | **5.60E-03** | 99 | **9.32E-04** |
| <u>Home large urban area scotland 20118 11</u> | 61 | **8.40E-03** | 138 | **8.88E-03** |
| <u>Whole body fat mass 23100 raw</u> | 62 | **9.76E-03** | 43 | **4.26E-07** |
| <u>Average total household income before tax 738</u> | 63 | **1.32E-02** | 271 | 2.20E-01 |
| <u>Age first had sexual intercourse 2139 raw</u> | 64 | **1.35E-02** | 39 | **1.31E-07** |
| <u>Age cataract diagnosed 4700 raw</u> | 68 | **2.23E-02** | 193 | **5.55E-02** |
| <u>North co-ordinate of birthplace in UK 129 raw</u> | 71 | **3.16E-02** | 81 | **1.73E-04** |
| <u>Skin colour 1717</u> | 72 | **3.42E-02** | 162 | **2.70E-02** |
| <u>Body mass index (BMI) 23104 raw</u> | 73 | **4.24E-02** | 52 | **3.55E-06** |
| <u>Birth weight of first child 2744</u> | 75 | **4.29E-02** | 342 | 3.84E-01 |
| <u>Malignant melanoma of skin C3 MELANOMA SKIN</u> | 76 | **4.31E-02** | 216 | **8.43E-02** |
| <u>Right intra-ocular pressure corneal-compensated 5254 raw</u> | 77 | **4.54E-02** | 72 | **6.98E-05** |
| <u>QRS duration during ECG 12340 raw</u> | 78 | **4.90E-02** | 194 | **5.68E-02** |
| <u>Longest period spent worried or anxious 20420 raw</u> | 176 | 3.73E-01 | 35 | **1.31E-08** |
| <u>Worrier/anxious feelings 1980</u> | 271 | 9.62E-01 | 70 | **4.80E-05** |
| <u>Neuroticism score 20127 raw</u> | 289 | 1.00E+00 | 69 | **3.65E-05** |
| <u>Never smoked 20116 0</u> | 293 | 1.00E+00 | 127 | **5.29E-03** |
| <u>Alcohol intake frequency 1558</u> | 290 | 1.00E+00 | 157 | **2.25E-02** |
| Genotype-phenotype (dbGAP) |  |  |  |  |
| <u>Body Height</u> | 1 | **2.61E-03** | 10 | **5.63E-08** |
| <u>Heart Function Tests</u> | 2 | **1.15E-02** | 36 | **1.23E-03** |
| Body Mass Index | 5 | 1.76E-01 | 1 | **1.13E-24** |
| Hip | 4 | <u>6.92E-02</u> | 7 | **3.65E-08** |
| Echocardiography | 6 | 2.11E-01 | 8 | **3.86E-08** |
| Body Weight | 7 | 3.14E-01 | 5 | **2.85E-11** |

<sup>1</sup>MacLean et al. 2019. Enrichment analysis performed with Inrichr. Traits significant in both studies are italicized and underlined, and $P_{Bonf} < 0.05$ are in bold and those trending underlined. All significant GWA catalog traits for this study's geneset were included here. For UK Biobank, a subset were selected as representative. The full analysis results are provided as supplementary tables.

Both canine behavior GWA genesets were significant for cardiovascular traits in all three human genetics datasets. Both were also significant for several other trait groups in the UK Biobank GWAS dataset: lung function, blood cell counts, skin color, behavior and demographics. Skin color associations are consistent with two of our behavioral loci which contain genes associated with coat color and pattern, but also expressed in the brain (*ASIP* and *MITF*). The brain traits significant in both genesets include some related to depression, demographics likely to be correlated with both socioeconomics and education, and reaction time. Six alcohol, tobacco and drug use traits were significant in MacLean’s geneset, but only one of those was nominally significant in our much smaller geneset. Two anxiety traits and neuroticism scores were significant in MacLean’s geneset, but not in ours.

We considered whether the comparative human GWA findings could be due to enrichment for body size/height. We asked if top human-GWAS traits associated with the canine genesets were also associated with height. Using the 162 complex-trait Polygenic Risk Scores (PRS) Atlas derived from UK Biobank GWA data^38^, we performed phenome scans in that population (Suppl. Text and Table S35). The traits with top associations with our geneset but not MacLean et al.’s – forced vital capacity and height – were extremely-highly significant for height PRS’s (both also for many anthropometric and diverse brain traits). In contrast, the top traits common to both genesets did not show that pattern: Distance between home and workplace was only significant, with Bonferroni adjustment, for the Years of schooling PRS; in contrast, the Height PRS ranked #72, *P*=0.44. Also, pulse wave arterial stiffness index (defined as height divided by systole peak-to-peak time) was significantly, and, by very far, most strongly associated with the PRS of Blood pressure (a known correlation^39^). This suggests the top traits shared by the two dog studies are not primarily due to their correlation with body size in humans.

### Analyses of hypothalamic single-cell expression data from juvenile mice

Based on data availability and tissue relevance^7^, we queried single cell RNA sequencing (scRNA-seq) data from hypothalamus (detailed in Suppl. Text)^40^. That dataset is based on dissection of a vertical column spanning the rostrocaudal span from preoptic to tuberal areas from pooled hypothalami of 2-4 week old mice of both sexes. We focused on the 62 neurons that were defined as clusters of sequences. First we identified the clusters depleted (*n*=3) or enriched (*n*=6) for both our GWA geneset and MacLean et al.’s by applying a 1-SD threshold to equal numbers of differentiation-ranked genes for each cluster (using rank-sum *P*). Hypergeometric testing with Bonferroni correction showed all three depleted clusters and enriched Cluster 61 were significantly depleted or enriched, respectively (Fig. 3B). We also tested the nine clusters against datasets of human genes associated with brain traits having approximately 700 or more genes. The traits were neuroticism and wellbeing (weakly powered), educational attainment, alcohol and tobacco use, brain structure and autism. Genesets were from GWA and gene-based GWA, except for the more speculative autism set based on exome sequencing and network-based predicted genes. We observed a general pattern whereby enriched clusters were strongly enriched for all the human datasets by comparison to the depleted clusters. Clusters 55 and 61 consistently had the strongest effects.

**Figure 3.**
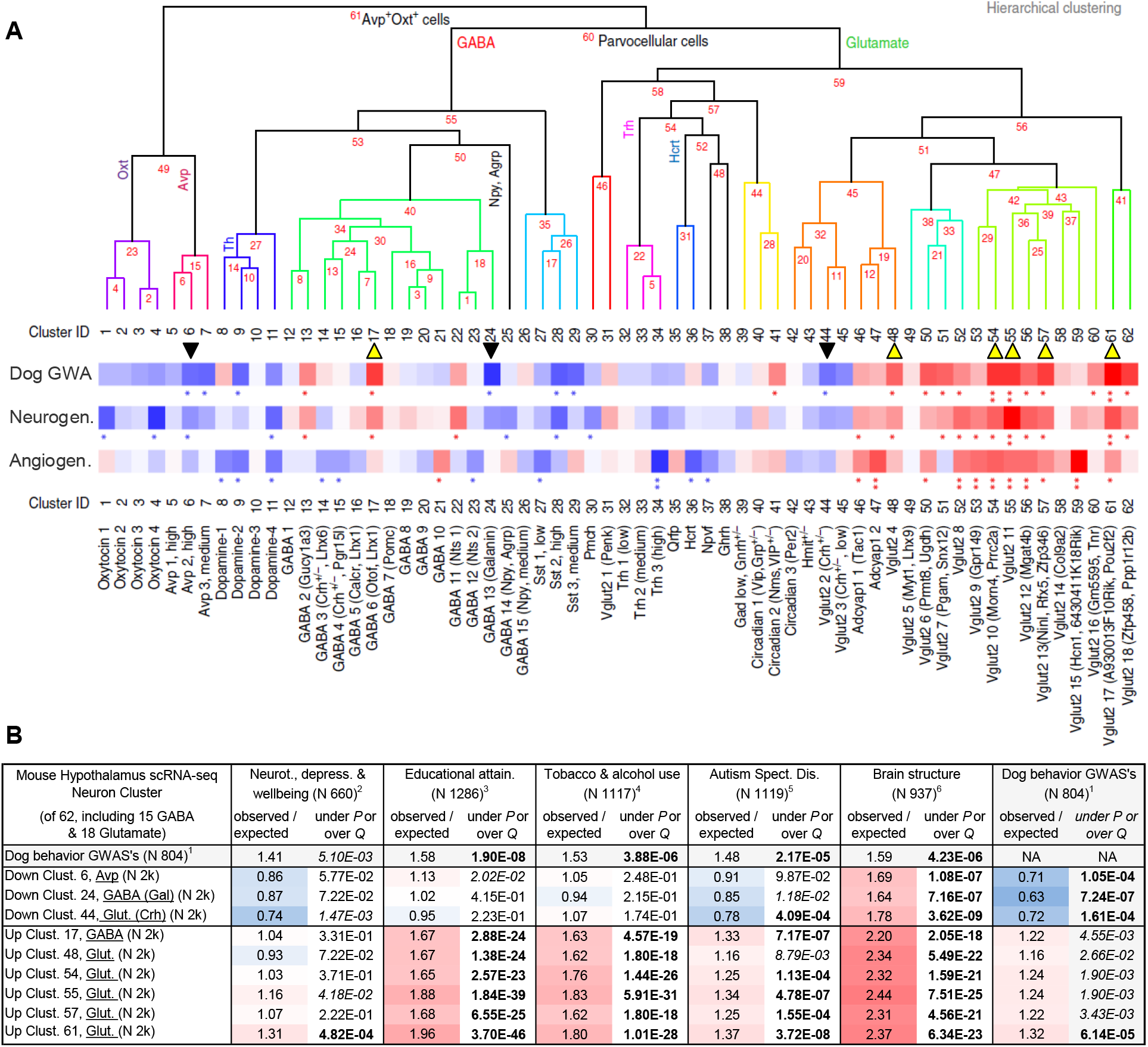
Relevance to specific neurons in the mouse hypothalamus and human neurogenetics. (**A**) Analysis of the canine behavioral GWA geneset vs. 62 neuron scRNA-seq clusters from mouse hypothalamus^40^, and GO genesets for neurogenesis and angiogenesis. The labeled cluster image was taken directly from Romanov et al. 2017 (ref. ^40^) and adapted. Yellow and black arrow heads mark the clusters depleted and enriched by 1SD from the mean in both the present and MacLean et al.’s (gene-based GWA corrected body mass)^6^ canine behavioral scans. Blue signifies depletion and red enrichment, and the number of asterisks corresponds to one or two SD from the mean for the combined canine geneset. (**B**) The marked depleted and enriched clusters in “A” were compared to the combined canine geneset and several brain-related genesets from human genetics. Note *P* and *Q* values are hypergeometric probabilities for depletion and enrichment, respectively; bold denotes significant with Bonferroni correction (*P*<8.47×10^−4^ threshold for 59 tests). Blue signifies depletion, red enrichment. Human datasets are provided in Supplementary Tables S39/40.

None of the six priority scRNA-seq clusters have been mapped^40^. Five of the six, including #55 and #61, are glutamate neurons. While few of the 62 clusters were reported to be sex-associated, #55 was predominantly male and #61 female^40^. We noted #55 is among the most differentiated clusters for the Y chromosome gene *Ddx3y*, and #61 is the top-most for the X-inactivation gene *Xist* (rank-sum *P*=6.19×10^−2^, *P*=8.38×10^−4^, respectively). Further studies are necessary to rule out other explanations for these sex effects (e.g., systematic dissection differences by age or sex). To begin to understand the biology of the six priority neurons, we compiled their differential expression of 155 signaling genes (Suppl. Text and Table S36). Among the findings relevant to our dog behavioral mapping and the question of pleiotropy with body size/mass and metabolism, *Igf1, Igf1r, Igf2r* and *Igfbp5* are differentiating for Cluster 55 (as are *Abcc8* and *Kcnj11*, indicative of glucose-sensing), and *Igf1r* is for #61. We next evaluated the most differentiating-genes shared by the six priority clusters compared to the rest of the 62 hypothalamic neuron clusters (Suppl. Text; Table S37). Those include many genes prominent in brain biology, such as *Spock2, Chrna4, Gabbr1, Kcnc1, Nrxn1, Celsr2, Megf8* and *Rbfox1*. We analyzed the top 18 most differentiated genes shared by the six clusters. 17 of 18 are associated with 374 human traits based on GWA. 12 of 18 are associated with 120 brain traits, most commonly in the area of drug use, followed by psychopathology and intelligence. Of the other six, one is associated with circulating IGF1 levels and the remaining five are highly expressed in the brain. Of the 18 top genes, seven are associated with height and six with BMI – with two overlapping. Four of 18 are associated with longevity (overlap of one each for height and BMI).

Transcription factors that significantly differentiate priority clusters supported the predictions based on both dog GWA genesets (Suppl. Text). For example, the top four Table 3 predictions – *POU6F1, POU2F1/2*, and *MEF2A* – were corroborated in the priority clusters. Out of the 62 hypothalamic neuron clusters, those four were differentiating in nine, four, seven and six clusters, respectively. Of the six priority clusters, all five glutamatergic clusters (but not GABAergic #17) were supported by at least one of those four transcription factors. All four were differentiating for Cluster 55 (as were the dog behavioral GWA genes *Pou6f2* and *Cux1*). Visual inspection of cluster differentiating genes revealed a neurogenesis and neurodevelopment profile for Cluster 55 (Suppl. Table S38). We thus tested all clusters for their content of dog behavioral GWA, neurogenesis and angiogenesis genes (Fig. 3A; Suppl. Text). This showed clear correlation between the mouse hypothalamic neuron cluster pattern and their overlap with dog GWA and mammalian neurogenesis genesets. Clusters 46-62 also correlated with angiogenesis. The restriction of *Lef1* and *Grin2b* expression to Clusters 51-62 suggests the implicated clusters are located more caudally, and are more immature and still undergoing synaptogenesis^41,42^.

### Testing for relevance to neurogenesis in human embryonic neocortex

We next tested for neurogenesis effects using open chromatin data from the developing neocortex of humans^43^. We found strong relevance of the combined canine GWASs for transcription factor motifs found in the neural precursor-enriched germinal zone *vs*. the neuron-enriched cortical plate (Fig. 4). That included transcription factors mapped and motifs predicted by the dog GWASs, and their paralogs. For example, the most strongly predicted transcription factor based on the dog mapping was POU6F1, which was supported by the mouse hypothalamus scRNA-seq data. It had the second strongest signal for germinal zone enrichment (*P*_*Adj*_=2.96×10^−92^). Two mapped dog genes, *CUX1* and *POU6F2*, were ranked eighth and eleventh, respectively (*P*_*Adj*_=3.23×10^−18^ and 5.09×10^−12^). The top neurogenic transcription factors in that study and those overlapping the dog GWASs are enriched for the Hox family (incl. POU, CUX and PAX subfamilies).

**Figure 4.**
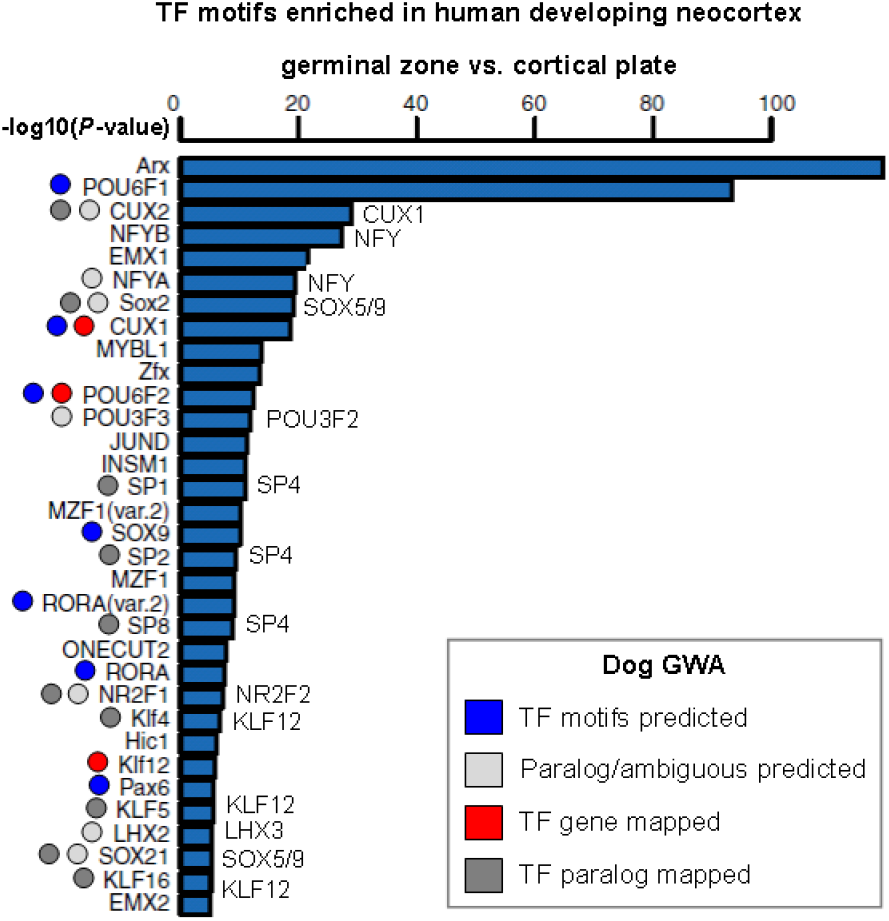
Relevance to neurogenesis in human embryonic neocortex. The combined canine behavioral GWA geneset from this study and MacLean et al.’s (gene-based GWA corrected body mass)^6^ was used to compare enriched transcription factor binding motifs (labeled on the right) to those enriched in developing neocortex germinal zone vs. cortical plate (labeled on the left). The figure was adapted from de la Torre-Ubieta et al.^43^.

### Detection of evolutionary footprints

We analyzed the genesets of the present and weight-corrected GWASs of dog behavior for relevance to selection (Table 5). Both were enriched for genes under positive selection in humans and domesticated dogs. Relevance of selection in 15 Chinese indigenous dog breeds hints that mapped variants predate breed creation and thus represent standing variation in wolves. Other domesticated animals are difficult to test because selection regions tend to be very large and contain many candidate genes, but the combined dog genesets were nominally significant for 666 candidate genes implicated in at least two of cattle, goats, pigs or sheep. Consistent with each other, both genesets are enriched for loss-of-function intolerant genes in humans^44^, and suggestive of depletion for genes with coding variation under selection in diverse wild and domesticated vertebrate species^45^. Both genesets are depleted for human single-trait associations and enriched for human accelerated-divergence regions^46^. In the preceding section we showed relevance to transcription factor motifs enriched in open chromatin in the germinal zone of developing human neocortex. That open chromatin is also enriched for human-gained enhancers, and their targeted genes are expressed in the progenitor-enriched laminae of the cortex^43^.

**Table 5.** Adaptive evolution and genomic demographics.

|  | Dog behavior GWA,<br>this study, n=108 |  | Mass-corrected dog<br>GWA<sup>2</sup>, n=715 |  | All genes, both<br>studies, n=804 |  |
| --- | --- | --- | --- | --- | --- | --- |
|  | Observed /<br>expected | under *P* or<br>over *Q* | Observed /<br>expected | under *P* or<br>over *Q* | Observed /<br>expected | under *P* or<br>over *Q* |
| Genesets compiled from the literature in this study<sup>1</sup> |  |  |  |  |  |  |
| Positive selection, dog breeds sampled in North Am. & Europe, n=1842 | 1.88 | **4.57E-04** | 1.35 | **1.00E-04** | 1.38 | **7.40E-06** |
| Positive selection, 15 Chinese indigenous dog breeds, n=963 | 1.11 | 4.20E-01 | 1.77 | **1.87E-08** | 1.64 | **4.01E-07** |
| Positive selection in at least two of cattle, goat, pig & sheep, n=666 | 1.61 | 1.07E-01 | 1.27 | 5.34E-02 | 1.3 | 2.88E-02 |
| Positive selection, human, n=1412 | 1.27 | 2.03E-01 | 1.38 | **3.03E-04** | 1.36 | **2.32E-04** |
| Loss of function intolerant, human, n=3154 | 1.28 | 6.02E-02 | 1.36 | **2.15E-08** | 1.34 | **2.42E-08** |
| Human accelerated divergence regions, n=1623 | 2.64 | **1.76E-08** | 1.85 | **2.49E-16** | 1.93 | **2.45E-21** |
| Protein-coding positive selection, vertebrates, n=550 | 0.433 | 1.54E-01 | 0.85 | 2.22E-01 | 0.785 | 1.05E-01 |
| Differential brain gene expression in 3 wild v. domest. mamm., n=121 | 1.02 | 6.41E-01 | 1.34 | 2.30E-01 | 1.32 | 2.24E-01 |
| Disease associated genes (ClinVar), n=3047 | 1.17 | 1.85E-01 | 0.87 | 2.82E-02 | 0.89 | 3.52E-02 |
| Haploinsufficient disease genes (ClinGen), n=294 | 3.65 | **8.14E-04** | 1.04 | 4.69E-01 | 1.31 | 1.09E-01 |
| Nearest gene to GWAS peaks (MacArthur Lab), n=6288 | 0.97 | 4.02E-01 | 1.03 | 2.21E-01 | 1.01 | 3.99E-01 |
| Single-trait pan-GWA meta-analysis (Watanabe et al.), n=1968 | 0.48 | 1.06E-02 | 0.62 | **1.97E-06** | 0.59 | **5.31E-08** |
| Top pleiotropic pan-GWA meta-analysis (Watanabe et al.), n=1968 | 1.27 | 1.43E-01 | 1.10 | 1.40E-01 | 1.10 | 1.23E-01 |

Hypergeometric *P* and *Q* are lower and upper cumulative distribution functions. Bold surpass Bonferroni significance for 26 tests, $1.92 \times 10^{-03}$ ; italicized surpass unadjusted threshold of 0.05.
<sup>1</sup>Gene lists and references provided in Supplementary Tables S39/40.
<sup>2</sup>MacLean et al. 2019.

## Discussion

Our canine investigations of behavior, genetics, brain imaging and comparative genomics are core facets of the neurobiology and evolution approach begun by Darwin and now referred to as personality neuroscience^47^. By mapping across dog breeds in three separate cohorts, we implicated 127 genes at 90 loci for risk of diverse normal and problem behaviors. We mitigated population structure and latent variables in the association analysis by using multiple cohorts with different breed makeup. Many of our findings are quasi-replicated: supported by the same or related traits being mapped in another cohort here or in prior studies^6,7^. In the Supplementary Text, we discuss the strengths and weaknesses of canine interbreed genome scanning. The findings are supported by our unpublished study of pedigree and mixed breed dogs with individual-level genotype and C-BARQ phenotype data^18^. Further support comes from the geneset overlap between the combined interbreed C-BARQ studies and gene-based canine cognition GWASs using individual-level genotype and cognitive testing data^17^ (hypergeometric *P*=2.13×10^−18^). While we can’t rule out the possibility of some false-positive discovery, the biological relevance of our GWA geneset suggests most loci are likely valid – even those mapped in a single cohort (incl. behaviorally-relevant genes *SHISA6* and *SMOC2*; Suppl. Text).

This work takes the first step to move canine behavioral genetics into the realm of brain physiology. We showed several of the loci presently mapped in multiple cohorts in this study are associated with brain structure differences previously detected across dog breeds^19^. That is supported by geneset enrichment for human GWA genes mapped for differences in brain structure – which includes six priority genes mapped in multiple cohorts here. While geneset analyses ranked limbic regions highest, our brain structure association findings and deeper results of geneset analysis showed diverse other brain regions were also significant (incl. thalamic and cortical). In contrast to humans^48^, complex genetic variations of dogs frequently have large effect sizes and thus have direct clinical and experimental utility^6,7,13^.

It is widely accepted that domestication is dependent on reduced fear, and that this occurred in dogs^49^. It is also clear that a high range of body sizes was selected for during the creation of dog breeds. A great extent of haplotype sharing across breed groups is attributed to selection for morphological traits such as body size and coat color^50^. Historical documentation also describes breed admixture for selection of temperament in breed creation. However the genetic evidence of that is limited and confounded by correlations of behavioral and morphological traits^6,7^. Coat color and decreased sizes of brain and body are the most commonly reported traits in matched domesticated vs. wild animals, but they are not universal^51^. Belyaev’s selection for tameness in foxes did not affect brain and body sizes. In contrast, Jensen and colleagues’ eight-generation selection for tameness in Red Junglefowl – ancestors of domesticated chickens – resulted in increased body size and smaller brain relative to body weight (^52^ and refs. within). There is thus strong evidence that selection for tameness does not necessarily affect body size, but that it can. Reduced fear was likely selected for in dog domestication, but it is unknown if body size was also selected. In contrast, breed creation had strong purposed-selection for diverse body sizes and behavioral traits^14,15,49,50^, and unintentionally selected for problem behaviors in smaller breeds^7,53-55^.

The opposite body-size biases of the two canine behavioral GWA genesets allowed us to stratify associated behaviors and biology (discussed in Suppl. Text). The two dog studies were supported by UK Biobank GWA relevance; and our phenome analyses allowed us to isolate body size effects and show the top traits shared by the two dog GWASs were unlikely to be due to their genetic correlations with human height. However, we cannot rule that out due to the different genetic architectures of body size in the two species, and the known association of growth signaling and blood pressure (incl. human *IGF1* variation^56^). Canine body size is known to be strongly correlated with behavior based on behavioral and genetic evidence^6,7,53-55^. For instance, analysis of 32,000 C-BARQ-phenotyped dogs from 82 breeds showed that behavioral clustering of breeds was explained more by body size than breed relatedness^55^. In humans, height is genetically weakly-correlated with brain traits including neuroticism, risk tolerance and smoking cessation^57-59^. As intracranial volume is correlated with body size in dogs^19^, it is with height in humans (phenotypic *r*=0.55; genetic *r*=0.26)^60^. Human brain volume is also genetically correlated with intelligence (*r*_*g*_=0.24) and the PI3K growth signaling pathway was implicated for both these traits^60^. That supports the interpretation that correlations of canine behavior and body size are due to pleiotropy and not population structure (Suppl. Text). This is also supported by the significant overlap of mapped genes and behavioral relevance between our GWASs and MacLean et al.’s controlling for body size^6^.

While both genesets were associated most strongly and consistently with the subiculum, ours weakly favored ventral aspects and the body size-controlled geneset predominantly implicated dorsal. The next-most implicated areas were hypothalamus and amygdala in ours and striatum in the body size-controlled geneset. The unique associations of tobacco and alcohol traits with the size-controlled geneset could be biological support for striatum relevance, but both those findings could simply reflect the much higher power of that geneset. The top anatomical rankings for our geneset seem consistent with defense from threatening conspecifics and predators. We are not aware of human GWASs of those traits and did not detect signal for possibly overlapping traits such as anxiety. One UK Biobank trait uniquely associated with our geneset uncorrected for body size was household income, which is known to be associated with both genetics of intelligence and the impact of one’s height on others that affects socioeconomic status^61^. Although the income signal was based on only four unlinked genes and could thus be spurious, all four genes in humans are associated with both height and intelligence or educational attainment, and are highly pleiotropic (median 122 ± 35 associations and 40 ± 22 traits (SD)). It is interesting that limbic regions and striatum are most highly implicated by our dog behavioral mapping. However, it is also possible other regions have similar effects, but rank lower due to a reduced density of relevant cells. Further studies are necessary to determine how robust and meaningful our anatomical enrichment findings are.

We also found associations with reproduction. Only the geneset uncorrected for body mass was enriched for uterus expression. This is consistent with the strong association of cattle stature with human height genetics and uterus gene expression^62^, and with the bidirectional association of endometriosis and diverse psychiatric traits in women^63^. Body and litter sizes are strongly correlated in wolves and dogs, hinting the same genetic network could be involved in determining body, uterus and brain sizes and functions^64,65^. Mendelian Randomization studies in humans show causal relationships between neurobiological and puberty traits, and BMI^66,67^. Consistent with that, the higher-powered geneset of canine behavioral GWA corrected for body mass showed enrichment for puberty traits and BMI. In humans and wolves/dogs, reproductive traits are also correlated with both external pigmentation and longevity^64,65,68^. Body-wide analysis of gene expression for age of menarche GWA in humans was only significant for five brain tissues, three of which were prioritized in this work: anterior cingulate, hypothalamus and pituitary^67^. That study implicated 16 transcription factors. Of those, three were mapped for dog behavior (*ELF1, NR2C2, MSX1*) or a close paralog was (human NR2F1 and SMAD3 implicated; dog *NR2F2* and *SMAD2/4* mapped); and two were significantly enriched in binding site predictions for both canine genesets (GATA, MSX1).

In addition to human genetics, the canine behavioral GWASs with and without control for body size were similarly supported by diverse geneset analyses (Suppl. Text). Both were most strongly enriched for the biological pathways axon guidance and morphogenesis, and gene annotation for alternative splicing and disease mutation. Analyses of evolutionary and genetic demographics show, or for some suggest, enrichment of genes that are pleiotropic, loss of function-intolerant, and under positive selection – including in humans. More specifically, there is also enrichment for human regions of highly accelerated evolution and containing human-specific enhancers that are active in neural progenitors.

Neurogenesis was strongly implicated in our analyses of dog GWA genes in both scRNA-seq data from juvenile mouse hypothalamus and for open chromatin data from embryonic human neocortex. Several transcription factor genes prioritized in this work are also known to have roles in adult neurogenesis, such as *Pou6f1, Pou3f2* and *Pax6* (Refs. ^69-71^). Evidence for neurogenesis networks includes *Pax6*, which regulates the mapped gene *Cux1*, which in turn regulates the mapped gene *Pou6f2*^72,73^. There was strong overlap of neurogenesis and angiogenesis in the same hypothalamic scRNA-seq clusters, consistent with their intimate association^74^. That suggests a possible pathophysiological effect for our candidate behavioral gene *ANGPT1*. Among the possible mechanisms linking body size and neurodevelopment, Fox-family transcription factor binding sites were ranked near the top for the combined dog behavioral geneset. Fox-family members, including *FOXO3* and *FOXA2* nominated here, have key roles in growth/IGF1-signaling, longevity and neurogenesis^75^.

Our scRNA-seq findings are supported by studies that used human GWA genesets from diverse traits to identify relevant cells in scRNA-seq data^76^. For instance, educational attainment and schizophrenia showed a Spearman rank correlation of cell-type association of 0.94 because both were enriched for telencephalon projecting excitatory neurons out of the 39 cell types queried, and both associations were enriched for neurogenesis and synaptic processes genes. Other analyses in that work showed relevance of both excitatory and inhibitory neurons for those traits as well as bipolar disorder and BMI, whereas neural progenitors and neuroblasts were prioritized for intracranial volume (shown there to be associated with height) and depression, respectively. Those findings seem to stratify our enrichment analyses, which implicate the broadest definition of neurogenesis from neural stem cells to maturing neurons. In agreement with us, those investigators stated the shared biology of psychiatric and cognitive traits is consistent with a general psychopathology factor. However, they concluded this effect – and other neuron-type associations such as with BMI – is probably mediated at the level of shared cell types rather than genesets. We propose a more parsimonious interpretation (explained in Suppl. Text) is consistent with a heritable psychopathology factor^77,78^: that risk genes and their networks overlap across functional and structural brain traits. As in dogs, this network is associated with height^57-59,79^, BMI^79^ and lifespan^80^ in humans. However, the large effect sizes in dogs seem likely to have resulted from body size selection under domestication. Further studies are necessary to assess that in other animals and in human “self-domestication”.

In conclusion, genome scans of dog behavior implicate genes associated with human behavioral, cognitive and psychopathological traits. Both dog and human GWASs consistently and strongly implicate neurogenesis at, or near, the top of enriched biological pathways^6-8,25,57,59,81-86^. We propose that evolutionary adaptation of behavior in mammals, and probably vertebrates, preferentially targets a genetic network for neurogenesis and neurodevelopment throughout life. This could at least partially explain the molecular basis for a general genetic factor for human psychopathology that is correlated with personality and intelligence. Our findings suggest that network is integrated with energy metabolism, growth, longevity and reproduction. Accepting survival and reproduction are the overarching priorities of natural selection, we believe the core evolutionary biology in play here is the balance of energy metabolism^87^ and growth/development (Suppl. Text). In addition to several highly implicated transcription factors (incl. Fox and Hox subfamily members, NR2C2/F2, SMAD2/4 and MEF2A), there is supporting evidence to nominate mTOR and its regulation by the ancient LIN28/let-7 loop as a core mechanism of that pleiotropy (incl. growth factor signaling-PI3K-AKT upstream; Suppl. Text B.10). It is clear that connected and opposing traits must be balanced, and not difficult to imagine how tradeoffs could impact risks of psychiatric traits. While evolutionary selection and neurodevelopment have long been implicated in psychiatric genetics, we believe our integrated proposal to explain the “p” factor is novel (e.g., see review of “p” factor genetics^2^ and discussion of possible mechanisms^78^). The cross-species conservation revealed here hints how genomics, neurophysiology and behavior could provide a scaffold for top-down descriptive and dimensional approaches to understanding the mind^88^.

## Methods

### SNP datasets and phenotype data

Three previously published SNP datasets were used in this study. Since the breed average phenotype data was not available for all the breeds included in each of the datasets, only those for which phenotypes were available were kept. The Boyko et al. dataset contained 423 subjects from 37 breeds genotyped for ∼45,000 SNPs (Affymetrix v.2 Canine array)^15^. The Vaysse et al. dataset contained 444 subjects from 29 dog breeds genotyped for ∼175,000 SNPs (Illumina CanineHD array)^14^. The Hayward et al. dataset contained 2885 subjects from 50 dog breeds genotyped for ∼160,000 SNPs (custom Illumina Canine HD array)^13^. Datasets were considered as independent, although the Vaysse dataset and the Hayward dataset used in this study share 192 subjects.

For the behavioral phenotype data, we used previously published C-BARQ data^7,89^. This data includes scores for the following problematic behaviors: stranger directed aggression, dog directed aggression, owner directed aggression, dog rivalry, stranger oriented fear, dog oriented fear, nonsocial fear, touch sensitivity, separation-related anxiety, attachment and attention seeking, predatory chasing, excitability, energy, trainability, persistent barking, urination when left alone and escaping/roaming. For the additional traits of size and lifespan we used data compiled from several sources^14,15^, the database curated by Dr. Kelly Cassidy at Washington State University available at http://users.pullman.com/lostriver/longhome.htm and from breed specifications published by the American Kennel Club (AKC) http://www.akc.org/.

### Genome Wide Association Analysis and gene annotation

Preparation of datasets, calculation of allele frequencies for each cohort, and removal of subjects with missing phenotypes were carried out in PLINK v.1.07(ref. ^90^). To maximize reproducibility across different SNP platforms, no dataset trimming or LD clustering were performed on the SNP data. Association analysis were performed with GEMMA v.0.94.1 (ref. ^16^). Population structure was removed by using the centered relatedness matrix correction. Association tests were performed using the univariate linear mixed model using the likelihood ratio test. Genome wide significance was based on Bonferroni adjusted *P*-value thresholds rounded down to the lowest order of magnitude: Vaysse dataset, 1 × 10^−7^; Boyko dataset, 1 × 10^−5^; Hayward dataset, 1 × 10^−7^. The genomic inflation factor λ was calculated as the median χ^2^(1 degree of freedom) association statistic observed across SNPs divided by the expected median under the null distribution^91^. Manhattan plots were generated with SAS v.9.4 from GEMMA outputs. Genome-wide significant hits were mapped on the CanFam3.1 genome assembly, UCSC Genome Browser^92^. Gene annotation was evaluated using the Broad Institute improved annotation V.1 track hub^93^ and human assemblies hg19/39.

### Effect direction and variance contribution regression modelling

To estimate effect direction and to determine the amount of variability that could be attributed to each relevant loci, we performed a regression modeling using a stepwise selection method. Since all hits detected were included, we performed a clustering step using Eigen decomposition on the first two dimensions. This decomposition allowed us to select single markers to represent a cluster and remove all the correlated markers that would have caused collinearity issues in the model. Each cohort was evaluated independently. All this analyzes were performed on SAS v.9.4.

### Brain-structure genetic associations

T2-weighted magnetic resonance imaging (MRI) morphometry data was derived from combined transversely- and sagittally-acquired images (using a 3.0 T MRI unit)^19^. Gray matter differences across breeds and sexes were measured as the degree of warping of per-subject maps to align with group-specific templates normalized by total brain volume (which was strongly correlated with body size). Independent Component (IC) analysis was performed using multivariate, source-based morphometry (GIFT software package for MATLAB^94^)^19^. IC loadings for each individual were evaluated though regression modelling using a stepwise selection method against the same relevant loci list used to detect effect direction and variance contribution. Regression modeling was performed on SAS v.9.4.

### Hypergeometric *P* value calculation

Of the 19,320 official protein coding genes in humans^34^, we found 16,080 to have a mouse ortholog. We arbitrarily used 80% of the latter number – 12,864 – as a conservative estimate of the number of genes likely to be expressed in the brain or to be represented across different mammalian species and types of datasets used here. Calculations were performed on SAS v.9.4.

### Geneset enrichment analyses

Geneset enrichment analyses were performed with Bonferroni multiple testing corrections on the Enrichr and DAVID algorithm and data servers^35,36^. Custom datasets were created for some enrichment analyses as follows. The hypothalamic scRNA-seq dataset was comprised of the top 2,000 genes, according to rank-sup *P* value, for each of the 62 neuron clusters reported^40^. Human genetics and evolutionary selection/genomic demographic genesets in Figure 3B and Table 5 are provided with sources in Supplementary Tables S39, S40. Hypergeometric testing (previous section) was used to calculate *P* values for those, and Bonferroni multiple testing thresholds were provided.

## Acknowledgements

We thank Marc Kent for providing opportunistically-collected brain imaging data from dogs, as described in ref. ^19^. This work was made possible by high quality and publically available canine data. For that we thank the groups led by Adam Boyko^13,15^ and Evan MacLean and Noah Snyder-Macklerand^6^, and the consortium led by Matthew Webster^6,14^. We thank the dog owners who contributed with biological samples and those who contributed to the C-BARQ database of behavioral phenotypes. We thank Dr. Kim McBride for his technical advice for designing this study. We thank the Stanton Foundation (Cambridge, MA) and Elisabeth Allison for their support.

## Authors Contributions

C.E.A. and I.Z. designed the study. I.Z. led the design of, and executed, the canine genetic analyses. C.E.A. led the design of, and performed, the comparative genetics and bioinformatics analyses, and the biological interpretations. J.A.S. collected and processed the C-BARQ data, and provided the behavioral expertise and, and E.E.H. collected and processed the brain imaging data, and provided that expertise. All authors contributed to the interpretation of results. C.E.A. drafted the manuscript with major contributions from I.Z., and participation from J.A.S. and E.E.H. All authors approved the final manuscript version.

