## Supplementary information: text and figures for "Genome scans of dog behavior implicate a gene network underlying psychopathology in mammals, including humans"

Index

A. Supplementary Text: Expanded results, discussion, methods, references (single Suppl. Info. PDF file; this file)
B. Supplementary Tables (single Excel file)
   1. Canine behavioral trait definitions (C-BARQ)
   2. GWA results, Boyko et al. cohort
   3. GWA results, Vaysse et al. cohort
   4. GWA results, Hayward et al. cohort
   5. GWA results and gene annotation of three cohorts combined
   6. Biological annotation for loci quasireplicated here or in published studies
   7. This study’s canine behavioral GWA geneset
   8. Body mass-controlled, gene-based canine behavioral GWA geneset (MacLean et al. 2019)
   9. Combined canine behavioral GWA protein coding genes with human orthologs genesets, this study and MacLean et al. 2019 body mass-controlled, gene-based GWA
   10. Human HUGO HGNC Protein Coding Genes used for selection of dog orthologs
   11. Geneset analysis, this study: Biological pathway, human KEGG (Enrichr)
   12. Geneset analysis, body mass-controlled GWA geneset (MacLean et al. 2019): Biological pathway, human KEGG (Enrichr)
   13. Geneset analysis, this study: Biological pathway, Gene Ontology, GO (DAVID)
   14. Geneset analysis, body mass-controlled GWA geneset (MacLean et al. 2019): Biological pathway, Gene Ontology, GO (DAVID)
   15. Geneset analysis, this study: Annotation, Swiss-Prot keyword (DAVID)
   16. Geneset analysis, body mass-controlled GWA geneset (MacLean et al. 2019): Annotation, Swiss-Prot keyword (DAVID)
   17. Geneset analysis, this study: Annotation, Uniprot tissue up (DAVID)
   18. Geneset analysis, body mass-controlled GWA geneset (MacLean et al. 2019): Annotation, Uniprot tissue up (DAVID)
   19. Geneset analysis, this study: Enriched gene expression, human BioGPS (DAVID)
   20. Geneset analysis, body mass-controlled GWA geneset (MacLean et al. 2019): Enriched gene expression, human BioGPS (DAVID)
   21. Geneset analysis, this study: Brain region enriched gene expression, mouse Allen Brain Atlas (Enrichr)
   22. Geneset analysis, body mass-controlled GWA geneset (MacLean et al. 2019): Brain region enriched gene expression, mouse Allen Brain Atlas (Enrichr)
   23. Geneset analysis, this study: Expression changes in transcription factor knockout mice (Enrichr)
   24. Geneset analysis, body mass-controlled GWA geneset (MacLean et al. 2019): Expression changes in transcription factor knockout mice (Enrichr)
   25. Geneset analysis, this study: Enriched transcription factor binding sites (DAVID)
   26. Geneset analysis, body mass-controlled GWA geneset (MacLean et al. 2019): Enriched transcription factor binding sites (DAVID)
   27. Geneset analysis, this study: Human GWAS Catalog (NHGRI-EBI)
   28. Geneset analysis, body mass-controlled GWA geneset (MacLean et al. 2019): Human GWAS Catalog (NHGRI-EBI)
   29. Geneset analysis, this study: Human UK Biobank GWA
   30. Geneset analysis, body mass-controlled GWA geneset (MacLean et al. 2019): Human UK Biobank GWA
   31. Geneset analysis, this study: Human Genotype-Phenotype Database, dbGAP
   32. Geneset analysis, body mass-controlled GWA geneset (MacLean et al. 2019): Human Genotype-Phenotype Database, dbGAP
   33. Geneset analysis of MacLean et al. 2019 canine behavioral GWA vs. all human GWASs with n>50k, compiled and curated (Watanabe et al. 2019)
   34. Transcription Factor binding site predictions called by two algorithms using this study’s and MacLean et al. 2019 mapped genesets combined
   35. Human Polygenic Risk Atlas analysis of top canine behavior-associated GWASs
   36. Annotation of 155 select signaling genes showing differentiation in six priority mouse hypothalamic neuron scRNA-seq clusters
   37. Top 20 genes implicated by 6 prioritized hypothalamic neuron scRNA-seq clusters
   38. Sample of prominent neurogenesis genes in mouse hypothalamus scRNA-seq neuron Cluster 55
   39. Genesets of protein coding genes with human orthologs tested for biological relevance in Table 5 and Fig. 3B (sources & references given in Table S40)
       a. Test human genetics geneset: Neuroticism, depression and subjective wellbeing
       b. Test human genetics geneset: Educational attainment
       c. Test human genetics geneset: Tobacco and alcohol use
       d. Test human genetics geneset: Autism spectrum disorder
       e. Test human genetics geneset: Brain structure
       f. Test data: Positive selection, dog breeds sampled in North Am. & Europe
       g. Test data: Positive selection, 15 Chinese indigenous dog breeds
       h. Test data: Positive selection in at least two of cattle, goat, pig & sheep
       i. Test data: Positive selection, human
       j. Test data: Loss of function intolerant, human
       k. Test data: Human accelerated divergence regions
       l. Test data: Protein-coding positive selection, vertebrates
       m. Test data: Differential brain gene expression in 3 wild v. domest. mamm.
       n. Test data: Disease associated genes (ClinVar)
       o. Test data: Haploinsufficient disease genes (ClinGen)
       p. Test data: Nearest gene to GWAS peaks (MacArthur Lab)
       q. Test data: Single-trait pan-GWA meta-analysis (Watanabe et al.)
       r. Test data: Top pleiotropic pan-GWA meta-analysis (Watanabe et al.)
   40. Data sources and references for Table S39
C. Supplementary Figures (single Suppl. Info. PDF file; this file)

1. Lifespan GWA

2. Preliminary epistasis analysis of loci quasireplicated in this study (see Suppl. Text)

D. Supplementary Data (separate Excel file)

1. Human GWAS data for HUGO HGNC candidate protein-coding genes

**Supplementary text index**

A. Expanded results
1. Interbreed genome scanning of behavior: gene annotation of previously reported loci
2. Interbreed genome scanning of body size and lifespan
3. Brain expression analysis of *ASIP*
4. a. Canine behavioral GWA geneset analysis: Transcription factor binding sites

b. Canine behavioral GWA geneset analysis: Evaluation of UK Biobank GWA genes and pleiotropy

c. Canine behavioral GWA geneset analysis: Evaluation of all well-powered GWASs compiled and categorized

d. Canine behavioral GWA geneset analysis: Relevance to hypothalamus scRNA-seq data

e. Canine behavioral GWA geneset analysis: Relevance to aging-associated DNA methylation

5. Epistasis analysis of prioritized main-effect loci
B. Expanded discussion
1. Strengths and weaknesses of canine behavioral genetics
2. Overview of present mapping validity, interpretations and utility
3. Proposed way forward for canine personality neuroscience and comparison to humans
4. Behavioral-body size correlations in dogs and comparative human genetics
5. Note on distinction of longevity-body size correlation within and across dog breeds
6. Correlations of energy metabolism and reproduction with canine behavior and body mass
7. Evolutionary genetics of canine behavior
8. Genetic associations of canine behavior and brain structure
9. Neuroanatomical implications of the canine behavioral GWA geneset
10. Behavioral adaptation acts on neurogenesis and mTOR pathways, suggesting a basis for pleiotropy
11. Dog coat color gene agouti (*ASIP*) has behavioral effects suggestive of domestication
12. Implications of behavioral GWA loci containing multiple biologically-relevant candidate genes
13. Epistasis analysis of quasi-replicated loci
14. Conclusions
C. Expanded methods
1. Epistasis
D. Expanded references

**Supplementary text**

**A. Expanded results**

**A.1. Interbreed genome scanning of behavior: gene annotation of previously reported loci**

The chr10:8.1 Mb locus was associated with the following traits in all three cohorts: body size, lifespan, attachment, escaping, excitability and touch sensitivity. The traits associated in two cohorts were barking, chasing, nonsocial fear, owner-directed aggression, rivalry aggression, separation anxiety, stranger-directed aggression and separation-related urination. This locus is the largest *F_ST_* region in the dog genome and is known to be associated with traits including body size ^1-3^, ear morphology ^1,3,4^, boldness^1^ and behaviors related to fear and aggression^5^. The locus, centered between *MSRB3* and *HMGA2*, has at least two markers present in three combinations that are associated with different traits^1,6^. Both *MSRB3* and *HMGA2* have neuropsychiatric relevance: hippocampus volume association with an intronic marker (*P=*1.94 x 10^-19^) and a role in neurogenesis, respectively^7-10^. *MSRB3* is the enzyme that catalyzes the reduction of methionine sulfoxide to methionine and *HMGA2* is a developmentally important chromatin modifying protein.

The chr15 locus was associated with body size, excitability, owner directed aggression and separation related urination in three cohorts. Barking, dog fear, rivalry aggression and touch sensitivity were associated in two cohorts, and attachment, dog aggression, lifespan, nonsocial fear and separation anxiety in single cohorts. We previously reported this haplotype was associated with several fear and aggression traits^5^. The risk haplotype, which is known to have two variants within an intron of *IGF1*, accounts for the largest contribution to small size across all but giant dog breeds^1-3,6,11,12^. Mammalian growth and lifespan are controlled through brain *IGF1* receptors; and *IGF1* has many neuropsychiatric-relevant functions, including in neurogenesis, axon guidance and other aspects of nervous system development^13^. GWA of human subcortical brain structure showed common variation in *IGF1* is associated with the volumes of the amygdala and brainstem^14^.

Chr18 was associated with many fear and aggression traits in our prior study, in which it had five associations quasi-replicated in both cohorts^5^. Here, that chr18 locus was associated with dog fear, nonsocial fear, size, stranger aggression, stranger fear and touch sensitivity in three cohorts. Barking, dog aggression, owner directed aggression, separation anxiety and trainability were associated in two cohorts, and attachment, chasing excitability, lifespan and separation related urination in a single cohort. We previously determined that, in a subset of breeds, the chr18 behavioral haplotype includes the *FGF4* retrogene insertion that most commonly causes chondrodysplasia^5^. But many breeds have the ancestral haplotype predating the retrogene insertion and manifest the behavioral effects. That interpretation was confirmed here by 14 breeds lacking chondrodysplasia and the *FGF4* retrogene insertion^15^, but exhibiting the chr18 behavioral effects: Beagle, Chinese Shar Pei, English Springer Spaniel, Cavalier King Charles Spaniel, Eurasian, English Setter, Siberian Husky, Bernese Mountain dog, German Shorthaired Pointer, Border Collie, German Shepherd dog, Labrador Retriever, Newfoundland and Doberman Pinscher. The two genes flanking the peak chr18 marker – *GNAT3* and *CD36* – are expressed in at least one cell type (taste receptors) and in multiple brain regions (amygdala, hippocampus and hypothalamus) in common. *GNAT3* encodes Gustaducin alpha, the Gα subunit that transduces signals for ligand-bound taste receptors in the mammalian tongue. In mice, it is also expressed in the vomeronasal organ and transduces alarm pheromone and predator-derived kairomone signals in the Grueneberg ganglion^16,17^. It also has chemosensory roles in mammalian airways and the gastrointestinal tract^18^. The enigmatic CD36 protein has diverse and complex roles in many cell types throughout the body, including in redox stress, inflammation, immunity, energy metabolism, angiogenesis, neurovascular physiology and behavior^19^. In the brain, it is expressed in neurons, microglia and endothelial cells^20^. Known behavioral effects include nearly four-fold increased expression of *Gnat3* mRNA in the amygdala of mice subjected to a pain stimulus and increased anxiety and aggression levels in *Cd36^-/-^* mice^21,22^.

Both cohorts with genotypes available for the X chromosome showed a haplotype spanning chrX:102.1 Mb is associated with dog fear, separation anxiety and size in two cohorts, and barking, chasing, excitability, lifespan, nonsocial fear, stranger fear and touch sensitivity in single cohorts. This locus is known to be associated with body size^2,5^, skull morphology^2^, sociability^1,5^ and fear traits^5^, and was discussed in detail in our previous study^5^. In brief, we showed that haplotype phasing can be used to split a very large linkage disequilibrium (LD) block into one haplotype associated with increased body size and reduced fear, and another with increased sociability. Based on positional evidence and biological relevance, we nominated *IGSF1* and *HS6ST2* as the genes affected at the two loci, respectively. *IGSF1* was subsequently shown to have a coding variant in the large body size haplotype^23^. IGSF1 is a transmembrane protein predominantly expressed in the hypothalamus and pituitary. Loss of function mutations manifest as a syndrome that includes deficiencies in thyroid and growth hormones^24^.

**A.2. Interbreed genome scanning of body size and lifespan**

Several of the haplotypes we previously reported to be associated with canine fear and aggression traits were known to be associated with body size^5^. Here we expanded that work by including a third cohort of genotyped pedigree dogs, and including breed averages of diverse other behaviors, body weight and lifespan. We identified nine loci associated with breed weight and supported by multiple cohorts here or by at least one prior publication using a different cohort (Table 1, Suppl. Tables S1/2). Seven of those were associated with at least one behavioral trait and seven with lifespan (with overlap of 6). Others mapped average breed weight to the same chr18 peak marker near *GNAT3* and *CD36* that we did^6,25^. They then used whole genome sequencing data to impute another variant downstream and nominated a chondrodysplastic *FGF4* retrogene insertion^15^ to be the causative variation^6^. Another study using whole genome sequencing detected association of that chr18 *FGF4* retrogene with height but not with weight (*n*=255 dogs). The present findings support our previous description of the major chr18 behavioral haplotype as being the recipient to the retrogene insertion^5^. Most notably, many breeds that lack the dominant chondrodysplasia phenotype carry this risk haplotype (discussed in the following section). Our interpretation is that the ancestral haplotype is associated with small body size and many behaviors.

GWA of breed stereotypes of lifespan confirmed prior findings at chr7:46.8 Mb (*SMAD2*), chr10:7.8-8.4 Mb (all 3 cohorts; *HMGA2*; Suppl. Fig. S1A), chr3:91.1 Mb (*LCORL*), chr15:44.8 Mb (*IGF1*) chrX:84.1 Mb (*IRS4* and *ACSL4*) and chrX:102.9 Mb (*IGSF1*)^3,26,27^. The autosomal associations are with small body size and increased longevity, and, in the X chromosome, with large body size and decreased longevity. All but the chr7 locus were also associated with behavior. The 2.5Mb fine-mapped chrX:84.1 risk interval – referred to as chrX Locus 1 – is not only associated with large body size, but also with bulkiness^23^. Both *IRS4* and *ACSL4* were nominated to contribute to those effects at this locus, but the interval contains many other genes^23,27^. We found three markers in this locus to be associated with size in one cohort, and the peak SNP was also associated with lifespan and the behavioral trait energy. That marker lies within the *PAK3* gene, which is flanked by *CHRDL1* upstream, and *CAPN6* and *DCX* (which is the last gene in the risk interval) downstream. All these genes are implicated in neurogenesis or neuroplasticity, which is well-established for *PAK3* and *DCX*^28,29^. This may be notable regarding one of the mouse hypothalamus single cell RNA sequencing clusters^30^ suggested below to be particularly relevant to our mapped geneset. Neuron cluster 55 (of 62), which was reported to be male specific and we found to have a strong neurogenic profile, shows across-cluster differentiation for over a third of the genes in this interval: *NXT2*, *ACSL4*, *TMEM164*, *AMMECR1*, *PAK3* and *DCX* (rank sum *P*<0.05). There is no such Cluster 55-enrichment in the approximately 2.5Mb upstream (only *IL1RAPL2*, -2.7Mb; *MORC4*, -1.7Mb; *MID2*, -0.5Mb) or downstream (only *HTR2C*, +2.8Mb). A genome wide DNA methylation association study of age-by-sex interaction yielded two autosomal and 12 X chromosome loci^31^. Of those 14, seven were attributed to specific genes – two of which are in or near the chrX Locus 1 region (*TMEM164, P=*6.75x10^-9^, and *MID2*, *P=*3.15x10^-10^). These two sets of observations hint that the chrX Locus 1 region may have general roles in neurodevelopment and longevity, and this could involve sex effects.

We identified two other haplotypes associated with small body size, increased longevity and behavioral traits in at least two cohorts: on chr13/*ANGPT1*, 3 cohorts; and on chr18/*GNAT3-CD36*, as presented above. Our within-breed direct phasing at the chr13 locus showed evidence of the reported selective sweep (Suppl. Fig. S1B)^1^. The haplotype with the G allele of the peak SNP had a minimal overlap region across breeds of ~50 kb between *ANGPT1* and *RSPO2*. The single exception was the Irish Wolfhound, which had a much larger haplotype containing both those genes. Notably, *RSPO2* has a rough coat variant that is fixed in this breed^32^. The A allele haplotype had a minimal overlap interval of ~650 kb that spanned both *ANGPT1* and *RSPO2*. The exception was in the Jack Russell Terrier, which did not span *RSPO2*. This is consistent with this breed’s coat variation (smooth, broken or rough). Importantly, these haplotype patterns do not correlate with breed lifespans. Our interpretation is that the selective sweep in this region was the result of human selection for coat variation at *RSPO2*, but not for the *ANGPT1* lifespan trait. Correlation analysis of trait scores revealed that lifespan is strongly correlated with body size and several behavioral traits (Fig. 1C).

**A.3. Brain expression analysis of *ASIP***

*Asip* was first cloned in mice and those original studies included Northern blot analysis of skin, whole brain and several other tissues from FVB/N mice^33,34^. *Asip* was only detected in skin, in which it was abundantly expressed (specifically, it is expressed in hair follicles according to immunohistochemistry). However, because most mouse neuroscience (incl. BioGPS and Allen Mouse Brain Atlas resources used in this work) is based on the C57BL/6 strain, which is null for *Asip*, we could not find any publication or public data resource showing its expression in the brain. There are reports that *Asip* is expressed in whole brains of rabbits, cattle and quail, but we did not find any that described expression across brain regions^35-37^. Here we report new analyses of public resources that provide some detail of brain region expression in cattle and humans.

The cattle data showed expression in hypothalamus (age unspecified) and in many regions in fetuses: medulla, cerebellum, cerebral cortex, pons and spinal column (NCBI EST Database). The human microarray data is derived from two *ASIP* probes and RNA from high-precision microdissection of 168 brain sub-structures classified within 27 brain structures sampled in a total of 6 brain donors (Allen Brain Atlas^38^). Using an arbitrary z-score minimum of 1.96 for each probe, the subiculum was positive in four donors, the central nucleus of the amygdala was positive in three donors and 13 other sub-structures were positive in two individuals. Of those, the top mean z-scores, starting with the highest were for the lateral tuberal nucleus of the hypothalamus, subiculum, supraoptic nucleus of the hypothalamus, parahippocampal gyrus and perifornical nucleus of the hypothalamus. The brain structures with the most positive sub-structures detected in any individual were the hypothalamus (11), cerebellar cortex (6), myelencephalon (5), pontine tegmentum (5), mesencephalon (3) and amygdala (3). Consistent with expression in several areas of the hypothalamus and cerebellum, and the primacy of the subiculum, RNA sequencing data from 14 brain regions showed the highest *ASIP* levels in the hypothalamus (mean 0.23 Transcripts Per Million or TPM, N 121 individuals), followed by cerebellum (0.18 TPM, *n*=173) and hippocampus (0.17 TPM, *n*=123) (GTEx^39^).

**A.4a. Canine behavioral GWA geneset analysis: Transcription factor binding sites**

Approximately one hundred transcription factor binding sites were significantly enriched for our geneset (Suppl. Table S23). 13 of the top 20 were also ranked top-20 in the MacLean et al.^40^ geneset (Table 3; Suppl. Table S24). We tested the set of 26 transcription factors associated with the top 20 binding sites for enrichment of GO biological processes. The overall pattern of hits was related to developmental processes, both generic and organ specific, and included 11 genes associated with both generation and differentiation of neurons. The seven sites in our top 20, but not MacLean et al.’s were not enriched for body size and related terms, but were also associated with development – most consistently with neuron development and differentiation (PBX1, GFI1, CUX1, NKX2-5, FOXD1 and GATA2/3). The remaining transcription factor, FOXQ1, is not well characterized but has roles in growth factor/PI3K signaling, aging/senescence, is ubiquitously and abundantly expressed in the brain, and is implicated in neuritogenesis^41^.

**A.4b. Canine behavioral GWA geneset analysis: Evaluation of UK Biobank GWA genes and pleiotropy**

Both genesets were significant for cardiovascular traits in all three human genetics datasets. Both were also significant for several other trait groups in the UK Biobank GWAS dataset: lung function, blood cell counts, skin color, behavior and demographics. [In the “Human GWA relevance” section of the main article, we provided evidence from phenome analyses of the UK Biobank GWA data that the top lung trait, “forced vital capacity” is strongly associated with height variation, but blood pressure traits are not. Of our 21 behavioral loci quasi-replicated in this work, 11 are associated with human blood traits (Table 1 of main article).] Elsewhere in this work we provide supporting evidence for the behavioral associations for two external pigmentation genes, *ASIP* and *MITF*. For this study’s focus on behavior, the most intriguing finding is that both genesets highly-ranked distance between home and workplace (ranked 1 and 2), and this was supported by distance to nearest major road and others. Such socioeconomic phenomena are presumably associated with intelligence and, as a result, social, educational and employment traits. That is consistent with both genesets being associated with age at first intercourse. Curiously, household income was significant in our geneset but not MacLean’s. A reminder that population stratification could explain at least part of an effect is a geographical association with both genesets and map coordinates of birthplace.

Another brain trait in the UK Biobank GWA that is significant in both genesets is reaction time (ranked 51^st^ in our geneset and 12^th^ in MacLean et al.’s). Among the psychopathological traits associated with both genesets were longest period of unenthusiasm/disinterest (23, 10) and longest period of depression (52, 29). In contrast to depression traits, two anxiety traits – and neuroticism score, which includes both of those – were only significant in MacLean’s geneset (ranked 176, 271 and 289 vs. 35, 70 and 69). This could be due to lack of power for our geneset or to differences in associations between height and depression or anxiety. At least one trait associated with alcohol consumption was significant in both (26, 6) and several tobacco/drug use traits are significant in MacLean’s geneset (discussed in Section B.9).

We next determined whether top human GWAS traits associated with both canine genesets are also associated with height. We used the Polygenic Risk Scores (PRS) Atlas of 162 complex trait PRS derived from UK Biobank GWA data to perform phenome scans in that population^42^. We focused on the top two UK Biobank traits from different domains in our results for both genesets – socioeconomics implicating intelligence and cardiovascular – and pulmonary and height, which ranked much higher for our geneset (Table 4; Suppl. Table S29, 30, 35). For our top trait and MacLean et al.’s second, distance between home and workplace, only the Years of schooling PRS was significant after Bonferroni correction (370 SNP PRS, *n*=160,231, *β*=0.0102, SE=0.00246, raw *P*=3.82x10^-05^, *R^2^*=4.51x10^-05^). That was followed by Putamen volume, Neuroticism, Fasting glucose, Hip circumference, Subjective well-being, Hemoglobin concentration and Parents age at death (65-187 SNP PRS range; *P*=1.78x10^-3^ to 1.67x10^-2^). Height ranked 72^nd^ (*P*=0.438). Our second-ranked trait and MacLean et al.’s third, pulse wave arterial stiffness index, was significant for a dozen PRS, beginning with Blood pressure (*P*=1.04x10^-40^) and Height (*P*=2.75x10^-15^). The only PRS for a brain trait was Years of schooling, which ranked eighth (*P*=5.86x10^-06^). The traits forced vital capacity and height were significant for 37 PRS, beginning with Extreme height and Height (both PRS achieved minimum-possible *P* of 1.00x10^-314^ for both traits). Both traits were significant for many anthropometric and brain-related traits: including Years of schooling (ranked 5 and 8, respectively) and a related trait, Infant head circumference (11, 7) and related traits, and Bipolar disorder (28, 23). Height was also associated with Schizophrenia and was suggestive for many brain traits, beginning with multiple depression and neuroticism traits, Neo-openness to experience, multiple reaction time traits, Internalizing problems, Hippocampus and Thalamus volumes, PGC cross-disorder traits, multiple alcohol and tobacco traits, Chronotype and Neo-extraversion. These findings support a biological association of both canine behavior GWA genesets with years of schooling in humans – a proxy for intelligence. Because the human trait distance from home to workplace is significantly associated with years of schooling and not height, this suggests that trait association with two canine behavioral GWA genesets is not simply attributable to height variation. However, due to the dramatically different genetic architectures of body size/height in dogs and humans, that possibility cannot be ruled out at this time (see Section B.4).

**A.4c. Canine behavioral GWA geneset analysis: Evaluation of all well-powered GWASs compiled and categorized**

For a comparison to well-defined and somewhat standardized human GWAS data, we queried a dataset that compiled all well powered GWASs (*n*>50,000). We tested the MacLean et al. geneset^40^ for overlap with 24 trait categories, but focused on the 12 with at least that number of genes (*n*=715; Suppl. Table S3). Activities (incl. diverse behavioral traits), Environment, Cognitive and Reproduction showed significant enrichment (Psychiatric was suggestive), but only Activities (hypergeometric *P*=2.09x10^-05^) and Environment (hypergeometric *P*=3.93x10^-03^) surpassed the Bonferroni threshold. We omitted the category Social Interactions, which only had 379 genes, from the formal analysis, but it was suggestive (hypergeometric *P*=7.95x10^-3^). This work’s far-lower powered geneset was not significant for any category, but also had the strongest signal for Activities. Combining the two genesets improved the significance for all four of those trait domains.

**A.4d. Canine behavioral GWA geneset analysis: Relevance to hypothalamus scRNA-seq data**

We tested whether single cell RNA sequence (scRNA-seq) data could suggest biological relevance of dog behavior candidate geneset. Based on our prior and current implications of hypothalamic relevance and the availability of a high-quality and extensive scRNA-seq dataset^30^, we made that our focus. This scRNA-seq study revealed 62 clusters of neurons in the pooled hypothalami of mice of both sexes at postnatal ages of 2-4 weeks. Sexual maturation occurs at 3 weeks, +/-1 and a small number of clusters were purely or predominantly female or male. As explained above, the following analyses were done with only dog and mouse protein coding genes with official human orthologs^43^.

We first screened the 62 hypothalamus neuron clusters for overlap with our and MacLean’s genesets. Three clusters were at least one standard deviation below and six clusters were at least one above the median of all clusters. We used hypergeometric testing to compare those nine clusters for relevance to four gene lists derived from human GWASs representing psychopathology, intelligence and drug use traits, and an autism geneset derived from case-control exome sequencing and network-based prediction (Fig. 4B). That table includes testing of those for overlap with the two dog behavior GWA genesets combined. At least one of the six up clusters was significantly enriched for all four human GWASs (#61) and five were significant for the three GWASs with >1,100 genes; none of the down clusters were nominally significant. The strongest enrichments and *P*-values were for glutamate Clusters 55 and 61.

It is important to understand the hypothalamus analysis above is exploratory. If true, such associations could be due to biological relevance of those precise clusters/neurons. Alternatively, the priority clusters could have general properties shared by neurons in specific developmental stages or across brain regions. Both GABA and glutamate clusters are present in the three down and six up clusters analyzed. However, five of six up clusters are glutamate neurons. Clusters 55 and 61, which had the strongest associations above, are among the very few clusters that appear to be very predominantly male or female^30^. Cluster 61 is based on only 3 sequenced cells and thus is not statistically robust. Despite Cluster 55 being based on 12 sequenced cells, we leave open the possibility the sex effect resulted from a systematic difference in tissue dissection.

We evaluated cluster differentiation by gene-expression using the comparison of each cluster vs. the other 61 (rank-sum statistic^30^). One indication the priority clusters are indeed related to the behavioral loci mapped in dogs is the differentiated expression of top transcription binding factors implicated by enrichment of their binding sites in those genes (Table 3). Pou6f1, which ranked first or second in both GWA genesets, had significantly differentiated expression in 9/62 clusters, including the priority Clusters 55 and 57 (priority clusters #54 and #55 also showed differentiation for the closely related Pou6f2). POU2F1/2 and MEF2A were ranked second and third according to binding sites in the GWA geneset and all are significant and enriched differentiation transcripts in priority clusters: Pou2f1 in 4/62 clusters, including priority Cluster 55; Pou2f2 in 7/62 clusters, including priority Clusters 54, 55 and 61; and Mef2a in 6/62 clusters, including priority Clusters 55 and 57. Those and other top binding sites support all clusters except #17, and most-strongly implicate #55.

Visual inspection of Cluster 55-differentiating genes revealed a neurogenesis and neurodevelopment profile (Suppl. Table S38). We thus measured overlap of the top 2,000 differentiation genes for each of the 62 neuron clusters with the combined canine behavioral GWA genesets, and the Gene Ontology genesets “neurogenesis” and “angiogenesis” (Fig. 4B). This showed correlation of the published clustering pattern with the dog behavioral GWASs and neurogenesis, most strongly in underrepresented Clusters 1-11 vs. overrepresented Clusters 46-62. In addition, the latter group of clusters is strongly correlated with angiogenesis. The top most overrepresented clusters for both dog GWA and neurogenesis were #55 and #61, which had the strongest enrichment and *P*-values for all four comparisons of human mental traits and diseases in Figure 4A. Further studies are necessary to understand what this means for our canine behavioral GWA findings. One possibility is that, say, the diverse neuropeptide neurons of Clusters 1-11, 23-30 and 33-39 are relatively mature, but the GABA Clusters 12-22 and glutamate Clusters 46-62 are undergoing neurogenesis and development. If so, genes related to neurogenesis status could be at least part of the signal implicating the six priority clusters. This would be consistent with the top biological pathway enrichment of axon guidance for both dog behavioral GWA genesets.

Although clusters with sequence data from as few as six cells have been mapped within the hypothalamus, none of our six priority clusters have been mapped^30,44^. Supplementary Table S36 illustrates the potential biological implications of the six priority clusters by showing their differentiation for expression of 155 signaling genes. For example, among the neuropeptide genes are proenkephalin and gastrin releasing peptide in Cluster 48, PACAP in #48 and #55, oxytocin and neurophilin in #54, Crh and Npvf in #55, and neurotensin in #57. Cluster 55 is particularly rich for neuropeptide receptor genes, which include those for Npy, neuromedin, somatostatin, Qrf pyroglutamylated RFamide, neuropeptide FF, hypocretin, melanin concentrating hormone, and neurotensin. A total of 20 G Protein Coupled Receptors that are targets of FDA-approved drugs^45^ are differentiation genes in the six priority clusters: 5 serotonin receptors, 4 adrenaline, 4 muscarinic acetylcholine, 2 GABA and one each dopamine, adenosine, histamine and purine. Among neurotransmitter biosynthetic or catabolic genes, the cannabinoid pathway genes *Dagla* and *Faah*, respectively, are significant in #55 and #61; and the cannabinoid receptor 1 is significant in #48 and #57. *Igf1* and *Igf2r* are significant in Cluster 55, and *Igf1r* in #54, #55 and #61. Given the prominence of Cluster 55 in this study and its differentiation for so many signaling genes, it is interesting that it is the only one of our priority clusters differentiated for *Eif4enif1* (rank-sum *P*= 7.03x10^-03^) and both Pumilio 1 and 2 (*Pum1/2*, *P*=5.20x10^-02^ and 1.13x10^-02^). Those proteins are known to bind specific mRNAs in neural stem cells and prevent their translation to regulate subsequent neuronal specification upon their release and translation^46,47^. While Pumilio proteins are generally considered to be translational repressors, they also have known roles as activators – including for our mapped candidate *FOXP1*^48^.

In a hypothesis-building context, we determined what gene expression distinguishes the six priority clusters from the 56 others. 94 genes were shared by at least four priority clusters. We ranked those by number of significant clusters, followed by the combined differentiation (rank-sum) *P*-value for all six. The top gene, *Spock2*, was significant in all six priority clusters compared to two clusters in the remaining 56. We evaluated the 18 of the top-20 ranked genes which have human orthologs (Suppl. Table S37). Several of those are cell surface signaling proteins, including nicotinic cholinergic and beta-type GABA receptor subunits (*Chrna4* and *Gabbr1*); the voltage-gated channel *Kcnc1*; and the adhesion and signaling receptors neurexin 1, *Celsr2* and *Megf8*. Two of the 18 top genes are RNA binding proteins, including the neurodevelopmental regulator *Rbfox1* which is a transcriptome hub gene in autism^49^. We tested each gene for associations in the human GWAS Catalog. This was intended to reduce the ascertainment bias in gene annotations, but is limited to genes with common human variations detected in well powered GWASs. All but one gene had at least one association. The remaining 17 genes had a total of 374 associated traits. All traits implicated in the geneset analyses were well represented except for demographics. 12 genes were associated with a total of 120 brain traits and a 13^th^ almost achieved genomewide significance. They included psychopathology (36 traits/6 genes), drug and alcohol use (26/9), intelligence (21/6), personality (15/4), sleep and circadian (8/3), performance of skills and memory (4/3), cognitive decline and neurodegeneration (4/3), and brain metabolism and structure (3/2). The top categories of psychopathology were neuroticism (18/4) and schizophrenia (8/5).

As dog behavior variations are enriched for height, weight and body mass index (BMI), it is notable that 7 and 6 of 18 are associated with height and BMI, respectively (overlap of 2). 4 of 18 genes are associated with longevity (overlap of 1 ea. for height and BMI). Only one of 12 genes associated with behavior was not associated with height or BMI. Of the six genes not associated with behavior, all are highly expressed in brain according to genome wide resources (mouse mRNA in BioGPS and Allen Brain Atlas; mRNA/protein Human Protein Atlas). Among the genes not known to be associated with height, or BMI, *SPOCK2* is associated with circulating IGF1 levels^50,51^. Dog behavior variations also suggested pleiotropy with coat color/pattern. *PRKCE*, which is a differentiation gene for five of the priority clusters, is associated with human sunburn risk in addition to height, weight, BMI, pathological gambling, suicide risk, reaction time and other traits. *RREB1*, another differentiation gene for five clusters, is associated with baldness, height, birth weight, math ability and insomnia. These various findings show common gene-level pleiotropy of behavioral traits and other traits including body size, and suggest plausibility of the mapping results and nomination of specific hypothalamus neurons as relevant. The interpretation is not straightforward, but the biology of the genes most differentiated in the priority clusters seem to lend further support for our study’s behavioral relevance as well as its enrichment for pleiotropic genes (incl. body size and BMI).

**A.4e. Canine behavioral GWA geneset analysis: Relevance to aging-associated DNA methylation**

Wang et al. recently developed a genome wide DNA methylation loci set selected for synteny across humans, mice and dogs^52^. They analyzed human and dog whole-blood DNA methylation patterns at 7,942 methylation loci near genes and identified 394 gene loci associated with age in both species, as well as mice. The geneset showing increased methylation with age was enriched for anatomical development; synapse assembly and regulation; and neuroepithelial assembly and regulation. The geneset showing decreased methylation with age was enriched for leukocyte differentiation and metabolic pathways. We tested whether those genesets are suggestive of relevance to the combined canine behavioral GWA geneset. We filtered the DNA methylation dataset to only include HUGO HGNC protein coding genes, n=7,690, of which 191 showed decreasing DNA methylation with age, 197 with increasing DNA methylation. 452 of the 7,690 genes were in our geneset of this study’s GWA combined with MacLean et al.’s (n=804 before filtering). We found genes with decreasing DNA methylation with age are suggestive of depletion, hypergeometric *P*=0.119 (population size 7,690; enrichment *P=*0.938) and those with increasing DNA methylation are suggestive of enrichment, estimated *P*=0.117 (depletion *P*=0.929). The overlapping genes with increasing methylation were ARHGAP28, COL12A1, RYR2, RYR3, SEMA6D, MAST4, MSX1, NR2F2, GDNF, HHIP, TBR1, TENM4, WNT5A, SNCAIP, ST8SIA2 and SYT1. Although neither was significant, the probability the two comparisons are different can be approximated, with some inflation, by combining the *P* values: *P*=1.39x10^-2^. This analysis suggests decreasing vs. increasing DNA methylation with age have opposite effects with regards to dog behavioral GWA genes. The caveats include the weak power for geneset enrichment analysis and the unknown relevance of whole blood epigenetics to brain function. If our findings were confirmed, it would suggest canine behavioral GWA genes are associated with increased DNA methylation with age. This hints at a plausible mechanistic link between behavior and aging-associated epigenetic-regulation of developmental and synaptic genes. That possibility is supported by both dog behavioral genes and human/mouse/dog genes with increased DNA methylation in aging sharing enrichment for developmental and synaptic genes.

In Table 2 in the main article, we reported that both our geneset and MacLean et al.’s are enriched for genes differentially expressed in WT mice vs. knockout of the Rett syndrome gene *MeCP2*, which is a methylated DNA binding protein involved in neurodevelopment. Two genes implicated in this analysis were also found in the *MeCP2* KO geneset, *TENM4* and *ST8SIA2*, both of which are known human psychiatric genes supported by mouse studies^53-57^.

**A.5. Epistasis analysis of prioritized main-effect loci**

Epistasis analysis is challenging due to computational burden. Here it is further complicated by having mapped 19 phenotypes (17 behavioral, lifespan and body mass), each done in three cohorts – which were genotyped in multiple platforms. We thus restricted our analysis to generalized pattern interpretation based on quasi-replicability across cohorts in this work (Fig. 1). That is, we conducted epistasis analysis of the main effect GWAS: chr10 (locus 2), 15, 18, 20 and 24 in all three cohorts, chr13 in the two cohorts in which it was detected, and chrX locus 2 in the two cohorts with chrX data. The pairwise interactions detected in all cohorts were chr10/24, 10/18 and 15/24. Of those tested in two cohorts, chrX interacted with 10, 13, 15 and 24. These findings are intended to be hypothesis-generating and should be interpreted with caution.

**B. Expanded discussion**

**B.1. Strengths and weaknesses of canine behavioral genetics**

Canine interbreed GWA has three outstanding advantages^1-3,5^. First, it allows mapping of breed averages of diverse phenotypes in unrelated cohorts with genotype data. Secondly, it makes it possible to map haplotypes that are fixed in many breeds. The implications of this are great if one considers that the average dog breed has approximately a quarter of its genome fixed for segments of 100kb or more^58^. Heritabilities of the present behavioral traits are on the order of five-fold greater across breeds as compared to within-breed^40^. Thirdly, as LD independently breaks down on both sides of functional variation in interbreed GWA, this has the effect of fine mapping^1-3,5^. Because dogs are bred under domestication, they have dramatically increased positive selection for desired traits and relaxed negative selection of deleterious variation. As a consequence of that and the strong selection of diverse traits, dog breeds present an extremely powerful population for genetic and translational studies^25,59,60^. The challenge to using interbreed mapping is population structure.

**B.2. Overview of present mapping validity, interpretations and utility**

In previous studies and here we mitigated the challenges of cryptic relatedness and population structure by using linear mixed models (implemented in GEMMA) for GWA analyses and by mapping the same traits in separate cohorts with different breed makeups^5^. The post population structure correction inflation factor (*λ*) of all GWASs in this work had a narrow range and averaged 1.14, slightly above the 1.05-1.1 that is considered benign. This could be due to cryptic relatedness and population structure. While the linear mixed model approach used here is efficient and powerful, it is still being improved for control of type I error and power in admixed and multi-ancestry cohorts^61^. For example, when the information is available, environmental correlations with trait-associated genetics that are not reflected in models of genetic relatedness should be included as covariates. There is also evidence that the control of inflation suggested by *λ*=1.0 is due to the cancelling out of inflation and deflation effects due to SNPs with higher and lower allele frequency differences than typical, respectively^62^. This issue manifests with both population substructure and ancestry admixture. These latter aspects as well as environmental covariates are clearly relevant to both pedigree and mixed-breed dogs. For instance, dog source (e.g., pet shop vs. shelter), purpose (pet, competition or work) and social environment (adults, children, other dogs or other pet animals) show correlations with behavioral traits and genetic markers^63^. Here we have mitigated that potential for false positive discovery by using three cohorts with different breed make-ups (and, thus, different cryptic relatedness and population structure). In that way, we prioritized 21 loci that were mapped for the same or a related trait in two cohorts, one here and the second here or in a published study.

Further support for our mapping comes from MacLean et al.’s similar GWASs^40^, but controlling for body mass and further correcting for inflation^64^. Of the 108 candidate protein coding genes we implicated at 90 loci, 19 genes were identified by MacLean et al. using gene-based mapping (hypergeometric *P*=6.91x10^-6^). While individual mapped loci cannot be assumed to be true due the caveats above, the full geneset shows strong biological relevance across species. Those findings suggest type I error is not a major problem here. We have also found support for the present mapping in an unpublished study^63^. There we report association analyses for 13 prioritized loci from this and our prior work^5^. For that, we created a 400-dog cohort with individual-level genotype and C-BARQ behavioral phenotype. It represented a community sample with roughly equal numbers of mixed-breed and pedigree dogs, and approximately one quarter of dogs had a clinical behavioral diagnosis. Between three models applied to the data, the results provided support for all 13 loci tested. We think of that work as a second exploratory phase of the present studies. Eventually, confirmatory (as well as new discovery) studies will require greater power. Genetically modified animal models will also be instrumental to validate and understand these findings.

Many of the geneset analysis results are consistent with behavior. This seems unlikely to be due to chance or to false positives resulting from body-mass selection that is unrelated to behavior: our geneset is small (*n*=108) and some loci have more than one gene positionally-implicated. Also supporting the mapping, many of the geneset analyses results are shared with those of MacLean et al.’s similar study, but correcting for body mass in the GWA (*n*=715). For instance, both genesets showed #1 enrichment rankings for brain in tissue-annotation and axon guidance for biological pathways in those respective analyses. They also shared significant enrichment for several behavioral and cognitive traits in the UK Biobank GWA analysis. Although we cannot rule out type I error due to population structure, these findings indicate this is not a major problem here. We prioritized 21 of the 90 the mapped loci based on quasi-replication for the same or related traits in two cohorts here or in published studies (Table 1). However, the geneset analyses suggest even loci mapped in a single cohort are true. For instance, *SHISA6* is known to sequester AMPA glutamate receptors in mouse hippocampus thereby inhibiting their desensitization during synaptic activity^65,66^. Another promising candidate is *SMOC2*, for which one haplotype (with a retroposon-mediated variant resulting in mis-splicing) is already known to be strongly associated with brachycephaly and possibly reproductive traits in breeds in which they are breed characteristics^67,68^. Human *SMOC2* variation is associated with various craniofacial/dental defects, and implicated in psychiatric, reproductive, pulmonary and dermatological (incl. pigmentation) traits^69-75^. There is also evidence human *SMOC2* has epigenetic states associated with brain structure and function^76^. Here we found the brachycephaly allele of canine *SMOC2* was extremely-strongly associated with urination when the owner leaves. This association was detected in the Hayward et al. cohort, which was the largest in this work and had strong representation of brachycephalic breeds. Eleven markers were significant for urination, and that peak marker was the single one associated (much less strongly) with chasing, fear of unfamiliar dogs, trainability and owner-directed aggression. Bladder^77^ (see mouse tissue mRNA expression in BioGPS^78^) and several brain regions involved in control of urination express *SMOC2*, including dorsal anterior cingulate (midcingulate; Allen Brain Atlas, mouse^38^). Of the three brain circuits controlling bladder function, the one associated with emotional aspects is centered on the midcingulate^79^.

A great limitation of GWA in general is the extreme delay in subsequent identification of the functional genes and variations, and thus their molecular mechanisms. In contrast, the geneset analyses conducted here suggest several handles for experimental testing. Our geneset is the result of GWA for many behavioral traits and is small. Unless the traits were genetically related, one would not expect the full geneset to be significantly enriched for biological relevance. We believe two factors could have contributed to our success in that respect. First, if the mapped functional variations and selection were old, then our positional precision for nominating candidate genes would increase. Second, existence of a core genetic network that is a common target of behavioral adaptive evolution could mean that many traits are associated with each other. Our prior and present results are consistent with both of those^5^.

Whereas the geneset analyses are exploratory, the convergence of multiple sources of supporting evidence suggests they have some basis in truth. For example, both canine genesets – with and without controlling for body mass – shared 13 of the 20 top most enriched transcription factor binding sites and those results were relevant to our mouse hypothalamus scRNA-seq analysis. We nominated six of those 62 scRNA-seq clusters based on their gene enrichment in both genesets. We subsequently determined the top three transcription factors implicated by their binding sites in the geneset enrichment analysis are *themselves* expressed and differentiating for those clusters. Thus, dog behavioral GWA genesets i) implicate a set of transcription factors because the member genes are enriched for their binding sites, ii) implicate several scRNA-seq clusters that specify individual hypothalamic neuron types in mice, and iii) those two findings are linked by the expression of the top three-ranked transcription factors in “i” being expressed in the implicated neurons in “ii”. Furthermore, the nominated clusters were preferentially and significantly enriched for genes from human GWASs of intelligence, psychopathology, drug use traits and brain structure. Another example is that neurogenesis (a top biological pathway implicated in diverse human personality and psychopathology GWASs^80-86^) and the subiculum of the hippocampus are strongly implicated by both genesets. Recent studies showed the subiculum to be a major site of adult neurogenesis in humans^87^.

**B.3. Proposed way forward for canine personality neuroscience and comparison to humans**

The best approaches for dog personality neuroscience and the comparison to human investigation are open questions^88^. Assuming similar studies across human populations were feasible, they would carry tremendous challenges that include ethical and legal constraints, and extremely high complexities of genetic/trait heterogeneity, environmental effects and socioeconomics^89-91^. Darwin’s first chapter of Origin of Species made a strong case that selection under domestication presented the best system for understanding evolution. Dog models have exceptional strengths that stem from that, including hundreds of isolated breed populations created in the evolutionarily-recent past^59,92-94^. Moreover, due to strong positive selection and relaxed negative selection, most complex traits are associated with a small number of variations of moderate to large effect sizes^1-3,5,25,95,96^.

The optimal way to understand canine biology and pathology is through a framework that includes epidemiology, behavioral and medical phenotypes, and genomics. However, it is not clear when such resources could materialize^92^. We believe interbreed mapping may be the most efficient discovery approach for understanding general dog and comparative behavior today^1-3,5,40^. Hayward, Boyko et al. compared individual-level body size measurements to breed averages for detection of genetic associations^25^. They made the surprising finding that, for the majority of mapped loci, breed stereotypes resulted in GWA *P*-values several orders of magnitude stronger than for individual body-size phenotypes. It is not clear why that is the case (the authors speculated it could be due to environmental effects), but it will be interesting to learn that as well as whether that is a general phenomenon. While interbreed mapping misses variations that are not common across breeds, it allows mapping of variations frequently fixed in many breeds. Paradoxically, this means both within-breed^97-100^ and across-breed^5,40,63^ approaches are necessary to fully understand each breed. We thus propose this and prior behavioral scans of breed-average traits to represent a first phase of discovery^5,40^, to be followed by a second phase using individual-level phenotypes in population samples of pedigree and mixed breed dogs, including dogs with veterinary behavioral diagnoses^63^. In addition to trait mapping by association, linkage or a combination^101^, individual-breed studies would allow for refined and diverse analyses of candidate risk alleles segregating within breeds.

Following the mapping of traits of complex genetics, there is a striking divergence of follow-on options for humans vs. dogs^92^. Due to negative selection, individual variations for human complex traits generally have minute effects^89^ and thus lack direct medical or experimental utility. In contrast, dogs have moderate-to-large effect variations associated with all of the diverse behavioral traits tested here, as has been reported for diverse other traits^1,2,25,95,96^. Prioritized variations can be validated and investigated in animal experiments or in veterinary or human translational studies. But the proposed approach also broadly-nominates genes, pathways, neurocircuits, mechanisms, etc.^96^. As is increasingly being done and we did here, those data can be analyzed together with diverse biological datasets to generate theories that are experimentally and translationally testable (Gilbert’s paradigm shift in biology^102^).

**B.4. Behavioral-body size correlations in dogs and comparative human genetics**

A major issue related to population structure is the well-established strong correlation of canine body size and diverse behaviors^5,40,103-106^. Here we will first present what is known about height genetics in humans and dogs, then discuss its correlation with behavior in the two species. The average male height in human populations varies from 137cm in the Mbutsi from Zaire to 183cm in the Netherlands: a 1.34-fold difference. The best-characterized large population with genetic data is the “white British”. Mean male and female heights in the UK Biobank are 175.7cm (6.7 SD) and 162.6cm (6.2 SD), and overall heritability is estimated to be *h^2^*≈0.8 (Ref. ^107^). Human height is one of the most polygenic traits studied to date^108^. It is estimated that human height is affected by almost 4% of common SNP alleles, or >100,000 SNPs with at least 1% minor allele frequency^109^. The top-contributing 1% of loci account for 6.5% of the estimated SNP heritability, or nearly four times less than the top 1% for the lowest-polygenicity traits like rheumatoid arthritis^108^. Based on effect-sizes of 20,000 SNPs, the polygenic score for the UK Biobank population only explains 40% of height variance^107^. [For considerations of population structure, polygenicity, pleiotropy and evolution in the understanding of human height genetics, see Ref. ^110^.] In contrast to a 1.34-fold height range in human populations, dog breeds have mean height and weight differences of 11.6- and 39.3-fold (male Chihuahua vs. Great Dane and Mastiff, respectively). Breed standards frequently include body size, but even different world populations of the same breed can vary in their statistics. A study of 93 breeds showed that seven loci explain 46-53% of body size variance for all breeds, and 64% for breeds with standard weights <41kg^12^. Another study of approximately 2,000 dogs from 158 breeds with both breed average and individual body mass and height phenotypes, used separately, showed 17 loci explained 80-88% of the variance in those traits^25^. A set of 330 village dogs in that study showed those variations only explained 30-40% of those traits, suggesting the 17 variants were more strongly selected in breeds. It was also observed that inbred pedigree dogs were smaller than their outbred members: in a breed with mean body mass of 20kg, an increase of inbreeding coefficient of 10% corresponded to a 1.2% body mass reduction. Thus, relative to human height, variation in dog height is very dramatically increased at the phenotypic level and very dramatically simplified at the genetic level. Genetic strengths include greatly reduced polygenicity and greatly increased effect sizes of variations. As noted above, complicating factors include frequent fixation of those variations in many breeds and the dramatically increased population structure in domesticated animals compared to humans.

Wilson et al. used C-BARQ behavioral phenotypes from 32,005 pedigree dogs from 82 breeds (*n*>50 ea) to cluster them according to their C-BARQ behavioral profiles^111^. They found membership to behavioral clusters was more-strongly driven by body size than breed-relatedness. Such a phenomenon is consistent with human genetic studies showing associations of height with many brain traits, including behavioral, cognitive and psychiatric^82,85,112^ and with human and canine brain imaging studies that show breed differences in brain structure^113^. Notably, tall stature and generalized overgrowth are risk factors for autism spectrum disorder^114^, but short stature and undergrowth are risk factors for schizophrenia^115^ (there is also evidence that head/brain growth rates, which are normally closely correlated with height, are abnormal in the first few years of life in both conditions^116^). That is consistent with the theory that autism and psychotic-affective conditions are diametric extremes of predominantly-normal variation in social and related mental states (vs. non-social and mechanistic)^117^.

We showed here that several dog body size loci are associated with canine brain structure differences. Our findings suggest loci associated with morphology frequently have strong effects on behavior and neuroanatomy. The chr15 allele we mapped for behavior is the same one known to be associated with canine small body size and increased lifespan^3,5,11^. The variation tagged by those genetic markers lies in the first intron of *IGF1* and results in reduction of its serum-levels^118^. The observation this locus has the most associations with different brain-structure IC’s is consistent with a body- and brain-wide hypomorphic effect. *IGF1*-modified animal models could be instrumental for dissecting its roles in brain development and behavior, and their correlations with organismal growth. Similarly, we identified candidate behavior and brain structure variation in genes highly expressed in brain, but predominantly known for their skin pigmentation biology. This refers to the secreted peptide ligand ASIP and the transcription factor MITF, but other related biology is apparent (e.g., transcription factor binding site enrichment analysis predicted POU3F2 to be relevant to the combined canine GWA geneset).

Human twin studies established that height and intelligence are associated, and that this is, at least in part, explained by individual variation in cortical volume and surface area^119^. Because those correlations are stronger than for intracranial volume, it has been suggested height genetics are more closely associated with use-dependent differences in humans^120^. The correlation of body and brain metrics is relatively weak in humans vs. dogs^113,120,121^; we presume that is due to strong selection under domestication for dog body size^12^. At least four of our top priority loci/candidate genes were also shown to be associated with brain structure differences in humans, *MSRB3* and *HMGA2*, *MITF* and *IGF1*^14,122^. It was fortuitous that MacLean et al. recently published similar studies to ours, but removing the effect of weight in the association analysis. Comparison of our geneset to their gene-based GWA shows they strongly reduced the number of genes associated with height in humans. In our geneset, height ranked #1, #1 and #6 in dbGAP, human GWAS catalog and UK Biobank GWA analyses, respectively; but it ranked #10, #108 and #60 for MacLean et al.’s geneset. Despite that, MacLean et al.’s geneset is significant, with conservative multiple testing correction, for height (incl. both #10 and #60 above) and several related traits.

Several of our behavioral associations are with known body-size alleles. We interpret the geneset results to rule out the possibility that our findings are largely false positives resulting from body size-associated population structure. Although MacLean et al. removed much of the effect of body size and our gene overlap does not seem high, the two genesets are significantly more similar than expected by chance. There are many parallels in the results of the analyses of the two genesets that would otherwise seem highly improbable, especially considering that our geneset only has 108 genes. For instance, both genesets rank axon guidance first in the KEGG pathway analysis and subiculum most consistent in the tissue analysis. Of the top 20 transcription factor binding sites associated with our geneset, 13 are also top-20 in the MacLean et al. geneset.

The UK Biobank GWA trait “distance between home and workplace” ranks #1 and #2 for our and MacLean’s genesets, respectively. While this hints at the relationship between intelligence and socioeconomics, the same dataset analysis showed income to be significant for our geneset but not the weight-corrected GWA. An alternative explanation is that the strongest correlation is between height and capital, rather than height and intelligence. MacLean et al. corrected for body size in the GWA and their geneset ranked height #60 (Bonf. *P*=1.08x10^-5^) and distance to workplace #2 (Bonf. *P*=3.14x10^-50^). Our geneset ranked height #6 (Bonf. *P*=1.38x10^-6^) and distance to workplace #1, 1.45x10^-15^. These findings seem too highly-significant to be spurious. In contrast, while our small geneset was only marginally significant for income after multiplicity correction (Bonf. *P*=1.32x10^-2^, rank #63), the 6.6-fold larger MacLean geneset was negative (Bonf. *P*=2.20x10^-1^, rank #271). Despite the weak statistical signal in our geneset, the association seems quite plausible because height and income are strongly correlated in humans. If these observed patterns were true, they would hint the two canine genome scans stratified cognitive and behavioral networks: distance to work genes enriched in both and height and income preferentially enriched in ours.

Mendelian randomization analysis of the UK Biobank shows nearly half of the correlation of height and capital is genetic^123^. The traits evaluated were income, educational attainment, job skill class and deprivation index; and the strongest statistical effect was for income. The study investigators referred to the 47% genetic contribution as “direct” and suggested likely “indirect” contributions could be intelligence and positive societal discrimination, both of which are well supported by published research. The evidence we discuss here (e.g., that height genes/variants are associated with differences in brain structure in humans and dogs) suggests that the height-intelligence association is at least in part direct. Pedigree dogs are comprised of hundreds of isolated populations that exhibit diversifying selection for many traits, including body size and behavior^5,12,40^. Wolf body size is highly variable within and across wild populations, and recorded specimens range from 12-79.4 kg. In contrast the *mean* weights of the US populations of smallest- and largest-breed are 2.25 and 77.1 kg. This shows how humans exerted very strong diversifying selection of body size across the overall population of pedigree dogs. Previously-accepted evidence that height is under selection in humans is now in question^110^. Because polygenic scoring of height is largely based on non-genome wide significant SNPs, subtle biases due to population structure result in false signals of adaptation that are statistically highly-significant. Selection of human intelligence is complex and apparently variable across populations, at least in part due to its associations with reproduction^124,125^.

To summarize this section, by comparing our canine behavioral mapping to a similar study that corrected for body mass, we were able to isolate effects of body size, which is under strong selection in pedigree dogs. The two genesets significantly overlapped – providing cross-validation – and both were enriched for brain gene expression and behavioral relevance. That is consistent with our study mapping true behavioral associations rather than primarily being artifacts of morphology (mainly, body size and coat pattern) and population structure^5,40^. The genetic architecture, and extent or effect sizes of pleiotropy, of the same traits in humans are not likely to be similar due to selection under domestication in dogs. However, there is rapidly-growing evidence from diverse human GWASs that pleiotropy is very extensive^126,127^.

**B.5. Note on distinction of longevity-body size correlation within and across dog breeds**

We noted in the main article that breed averages of body weight/height and lifespan are negatively correlated across most, but not all, breeds; and the two traits are associated with the same genetic variations^3,27,128,129^. However, it should be noted that within breeds there is either no correlation or, consistent with diverse other species, there may be a positive correlation^130,131^. Evidence from single breeds suggests body mass index and excess consumption of food are important factors in longevity, but height is not^131^. Other factors are mixed breed status and neutering, both of which are associated with increased lifespan^132^.

**B.6. Correlations of energy metabolism and reproduction with canine behavior and body mass**

Whereas our geneset was strongly enriched for human height genetics, it and the geneset correcting for body mass were enriched for human BMI and related traits, and cardiovascular traits (Table 4). In the main article, we mention there is also evidence in this work that reproductive traits are also correlated, including uterus gene expression, litter size and puberty timing. Curiously, uterus was the only significant tissue transcriptome enriched in our geneset after multiplicity correction. In the main article, we note our uterus finding and the strong correlation of body and litter size in canine breeds^133^ suggest the effect is related to enrichment of body size variation in our geneset. Therefore, the biological effect is likely to include – but not be limited to – gene functions involved in whole body and uterus growth (incl. body-organ size scaling), and litter size is constrained by uterus size. In women, endometriosis and diverse psychiatric traits are significantly and bidirectionally associated^134^. The first-ranked and significant gene ontology pathway enriched in our geneset was embryo development, but it only ranked #184 for MacLean’s weight-controlled geneset (Bonf. *P*=1.00). This observation is consistent with developmental or growth pathways. Notably, maternal height is strongly associated with gestational duration and birth size due to maternal/uterine and fetal genetic effects^135^. The same variation, most of which is more clearly associated with birth size, is in turn associated with cardio-metabolic traits^136^ that are enriched in both canine behavioral GWA genesets.

These observations about reproduction are also interesting because its inhibition by stress is a major barrier to domestication, and mapping of fear/defense traits implicated the hypothalamus and pituitary here and in our previous study^5^. In the main article we mention associations of canine behavioral genes with UK Biobank GWA for puberty timing in both sexes. Environmental and body-wide signals that impact reproduction are integrated in the hypothalamus, which, via the pituitary, regulates gonadal function (referred to as the HPG axis). The hypothalamus-pituitary also i) regulate homeostasis and the stress response to the adrenal glands (HPA axis) by secreting corticotropin-releasing hormone and adrenocorticotropic hormone, respectively; ii) regulate metabolism by secreting thyrotropin-releasing hormone and thyrotropin, respectively; and iii) regulate body size by secretion of growth hormone-releasing hormone and growth hormone, respectively, resulting in IGF1 secretion by the liver. Hypothalamic nuclei are not only connected indirectly, but extensively interconnected directly. It does not seem surprising these aspects of physiology could be genetically integrated.

**B.7. Evolutionary genetics of canine behavior**

The two GWA genesets for dog behavior– with and without correction for body mass – were similarly enriched for several evolutionary footprints, but differently for others. Both genesets were significant for genes positively selected in breeds sampled in North America and Europe. However, only the weight-corrected GWA geneset was significantly enriched in the half-as well powered list of genes under selection in 15 Chinese indigenous dog breeds. The relative enrichment of the two GWA genesets for North American/European-sampled breeds vs. Chinese breeds suggests the two breed samples differ in the primary variation we mapped in our behavioral GWASs (i.e., enriched for body-mass variation). In contrast, the weight-controlled behavioral GWASs implicate genes under selection in both breed samples, but more so in the Chinese breeds. If shared genes are on the same haplotypes, then this would indicate those variations predate the divergence of those two breed samples.

There are abundant reports of genomic regions under selection in domesticated cattle, pigs, sheep and goats, but these are generally very large and contain many genes. To avoid small signal-to-noise in such gene data, we compiled a relatively small list of genes implicated in at least two of those species (n=666). Enrichment for that domestication gene list was marginally significant without multiplicity correction in the MacLean geneset and in the two genesets combined. The fold-enrichment for human positive selection genes (*n*=1412) was similar to domesticated mammals, but the MacLean and combined genesets were significant after multiple testing correction.

Dog GWA genes were depleted for selection of protein-coding variation in vertebrates and, consistently, with enrichment for loss of function-intolerant genes in humans. Possibilities for the basis for such constraint include protein structure, high numbers of interaction partners, hub-status in a network, and critical or highly-optimized protein function. Among the strongest statistical signals in the evolutionary analyses was enrichment for human accelerated-divergence regions. That is consistent with our finding that the dog genesets and the transcription factors predicted to regulate those genes are i) overrepresented for the top transcriptional motifs enriched in open chromatin in the neocortical germinal zone of human embryos; and ii) that open chromatin is enriched for neural progenitor-active enhancers gained in humans^137^.

**B.8. Genetic associations of canine behavior and brain structure**

We showed here that several dog body size and two coat-pattern haplotypes are associated with variation in canine behavior and brain structure. For example, the mapped *IGF1* allele is also associated with small body size and extended lifespan. Our finding this allele is associated with several canine brain structure IC’s is consistent with the fact this allele results in reduction of IGF1 serum-levels^118^ and the possibility of a body-wide hypomorphic effect. Regarding coat genes, we found behavior/brain structure-associated variations implicating two genes highly expressed in brain, but predominantly understood for their roles in external pigmentation: the secreted peptide ligand Agouti-signaling peptide (*ASIP*) and the transcription factor *MITF*. Strikingly, six of our top-priority candidate genes at three loci quasi-replicated for behavior (two also associated with dog size and one coat pattern) have also been reported to be associated with differences in both brain structure and function in humans: *MSRB3* and *HMGA2*, *GNAT3*, *MITF* and *FOXP1*, and *IGF1*^7,14,122,138-140^.

The six IC’s are classified according to the brain structures involved and named for their suggested functions of those. IC4, called “Fear, stress and anxiety: HPA axis” based on the brain structures affected, is approximately tied with IC2 and IC3 for the most marker associations. The chr10, chr15, and chr18 loci are associated with many traits and several IC brain networks, suggesting broad or dispersed anatomical effects. In contrast chr13, chr20 and chr24 allow for pairwise consideration of mapped behaviors and brain networks. In the following, we italicize mapped traits that could hint at testable hypotheses. IC2, “Movement, eye movement and spatial navigation” regional network was associated with the chr15 locus mapped for *escaping* and owner-directed aggression in three cohorts, and *chasing*, separation anxiety and emotional urination in single cohorts. IC3, “Social action and interaction”, was associated with the chr18 locus mapped for *excitability* and nonsocial fear in two cohorts, and *aggression directed at unfamiliar dogs*, *separation anxiety* and *attachment* in single cohorts. IC5, “Olfaction and vision”, was most strongly associated with the chr24 locus mapped for *nonsocial fear*, *aggression directed at familiar dogs* and separation anxiety in two cohorts, and *chasing*, energy, *owner-directed aggression*, touch sensitivity and urination in single cohorts.

**B.9. Neuroanatomical implications of the canine behavioral GWA geneset**

Human and dog GWASs of behavior, cognition and psychopathology have implicated diverse brain regions^5,40,80-86,141-143^. The two dog mapping studies considered here – with and without correction for body mass – supported each other by implicating the subiculum most strongly (Table 2; Suppl. Table S21, 22). The subiculum, which receives the main output from the CA1 field but has extensive other connections, is involved in memory, cognition, physiological and emotional stress, and motivation^144^. However, there were also differences between the two dog studies that hint at stratification of traits and genetic pathways associated with body size. For instance, MacLean’s dataset was predominantly enriched for dorsal vs. ventral subiculum, whereas ours weakly favored ventral. After subiculum, both genesets were most strongly enriched for different brain regions with prominent connections with the subiculum. Our geneset was next most enriched for hypothalamus (incl. preoptic and ventromedial areas), followed by amygdala and associated areas (bed nucleus of the stria terminalis, amygdalopiriform area, and suggestive for different parts of the medial amygdalar nucleus (posterodorsal most significantly, *P_Adj_*=5.59x10^-2^)). The much-higher powered geneset correcting for body size was next most enriched for all parts of the striatum; CA1 and the anterior cingulate area were also near the top, and many other areas were significant, including the posterodorsal medial amygdalar nucleus (*P_Adj_*=9.17x10^-3^) and several areas of the thalamus and cortex.

Our geneset’s enrichment for hypothalamus is consistent with pleiotropy of body size and behavior. Specifically, preoptic and ventromedial hypothalamic areas were most strongly implicated. That brings to mind the most important defense pathways across animals: responses to direct threat-stimuli (i.e., distinct from contextual cues) from the same species (MeA > MPO > VMH > PMM > PAG) and predators (MeA > VMH > PMM > PAG)^145^. Stratification of traits according to body size pleiotropy is supported by testing of our geneset’s 24 genes associated with hypothalamus, amygdala and BNST for UK BioBank GWA relevance (Enrichr^146^). The top two traits were height and birth weight (*P*/*P_Adj_*=1.98x10^-4^/1.70x10^-1^ and 2.93x10^-4^/1.26x10^-1^, respectively). The next two traits are related to those, but the fifth is the top trait shared by the two dog genesets – distance between home and workplace (*P*/*P_Adj_*=2.09x10^-3^/1.99x10^-1^; 5/11 genes for this trait were shared with genes for height trait) – which we showed by phenome analysis here is unlikely to be predominantly due to height. This interpretation is also supported by UK BioBank enrichment analysis of the 54 genes enriched for subiculum in one or both genesets. The top trait is the same as the top one shared for enrichment in both full genesets: distance between home and workplace (*P*/*P_Adj_*=6.00x10^-12^/5.14x10^-9^). By contrast, height ranked 65^th^ (*P*/*P_Adj_*=1.51x10^-2^/2.04x10^-1^). Our brain structure associations provide physiological support for stratification of behavior and genes pleiotropic for body size. Three loci were quasi-replicated for association to IC5, “Fear, Stress and Anxiety-HPA Axis” (correlated with structure differences in hypothalamus, hippocampus and amygdala): chr10, *MSRB3-HMGA2*; chr15, *IGF1*; and chr24, *ASIP*. The first two alleles are also associated with body size. The third allele is pleiotropic for coat color, but we reported here *ASIP* is expressed in the brain, and, in humans, most strongly in the subiculum, amygdala, hypothalamus and cerebellum.

By contrast, the striatum enrichment of MacLean’s geneset supports its strong associations with seven UK BioBank alcohol and tobacco GWA traits. Those were also enriched in our geneset, but only two were nominally significant and the one significant after Bonferroni adjustment is ambiguous (frequency of needing a morning drink after heavy drinking) and had a *P* value 20-times larger than for the other geneset. We believe these geneset differences are relative, not absolute. They seem likely to be due to our geneset’s greatly reduced power and enrichment for pleiotropic behavior/body size variations. It is important to understand that the genetic architecture of behavior and body size is very different in dogs and humans^1-3,5,11^. Several pleiotropic variations we mapped in dogs have large effect sizes for both behavior and body size, but such variants are presumably extremely-rare in humans. Compared to natural selection in mammals in general, human selection of dog size has exaggerated the effect levels of body size variation and its pleiotropy. Our ability to stratify such effects was greatly potentiated by that, and the fact our geneset was biased for the largest effect sizes (which are enriched for body size) and the other geneset was designed to exclude body size variation.

Those brain regions found most consistently at the top of our enrichment analyses are reminiscent of the limbic theory of emotions initiated by Broca, greatly developed by Papez and Jakob, extended by MacLean and Panksepp, and still undergoing regular study and critique^147,148^. With the exception of the anterior thalamic nucleus and the mammillary bodies of the hypothalamus, this work implicated every key node of that theory. The area most consistently nominated by both canine behavioral genesets was the subiculum, which projects, via the axonal superhighway called the fornix, to the mammillary bodies. We checked for non-significant enrichment of the two “missing” areas in both dog genesets. The mammillary bodies were very weakly and not even nominally-significantly enriched in our geneset, and were non-significantly depleted in MacLean’s. Parts of the anterior thalamic nucleus were weakly enriched in both, beginning with the anteromedial nucleus in both (ours nominally significant, MacLean’s non-significantly). It is an open question if these and other significantly-enriched areas of the brain could be important in behavioral adaptation as others we nominated here. That seems likely, but it is possible limbic regions are particularly central or somehow more malleable, whereas other brain regions have more subtle or distributed effects. There is active pursuit of these questions using brain imaging to identify associations with psychopathology and a major focus is the distinction between limbic and cortical anatomy^149-152^. For instance, a lesion-approach study showed the strongest statistical signals for an internalizing factor were in regions of interest in the basal ganglia and amygdala, and, for an externalizing factor, in the orbitofrontal cortex and hippocampus^153^.

It should be noted that our scRNA-seq analyses of mouse hypothalamus used a dataset which only included the rostrocaudal aspect from the preoptic to tuberal regions, and thus lacked the mammillary area. We noted in the main text that scRNA-seq neuron clusters enriched for neurogenesis and angiogenesis are presumably the most immature and could represent the caudal end of a rostro-caudal axis of development. Given mice of different ages and sexes were pooled for that scRNA-seq study, it seems possible variability in dissection could explain some of the male or female specific clusters. Such an effect would be key here because the top two implicated neuron-clusters according to canine behavioral mapping were specifically or predominantly male (#55) and female (#61). Put another way, if it were true those two clusters are the most immature and posterior, is it possible they resulted from age or sex differences in dissection of the posterior end of the hypothalamus (e.g., by inclusion of neurons from the medial mammillary area)? None of the scRNA-seq clusters prioritized here have been mapped anatomically, but such efforts have been initiated^44^. It will also be important to conduct scRNA-seq analyses of all brain regions. That will enable another phase of theory creation and testing on the way toward the understanding of behavior at the molecular and neurocircuits levels.

**B.10. Behavioral adaptation acts on neurogenesis and mTOR pathways, suggesting a basis for pleiotropy**

Our geneset analyses of canine behavioral GWASs suggest there is a genetic network that is preferentially targeted by behavioral adaptive evolution (see section B.7). We showed here the canine behavioral GWA genesets are also associated with postnatal neurogenesis in the hypothalamus scRNA-seq data (especially cluster 55) and open chromatin data from embryonic human neocortex^30,137^. We also showed the subset of loci quasi-replicated in this work is associated with differences in dog brain structure. Those findings converge with a broad body of knowledge^141^ and the recent explosion of human GWA evidence that human behavioral, psychopathological, cognitive, and drug use traits are strongly associated with neurogenesis/neural stem cells and neurodevelopment^5,40,80-86,142,143^.

The fact that canine body size loci are most strongly enriched for the IGF1 pathway^154^, which has major roles in neurogenesis^155^, supports the possibility that body size and behavior are functionally associated. That is also supported by human twin studies showing that intelligence and height are associated, and this is accounted for by individual differences in cortical volume and surface area^119^. Our mouse scRNA-seq evidence also shows angiogenesis is correlated with neurogenesis in the same cells. This is consistent with our mapping of the *ANGPT1* locus and geneset enrichment for cardiovascular traits. It is also supported by the fact that neuronal stem and progenitor cells in the adult hippocampus of mice maintain their niche by secreting vascular growth factors^156^. *ANGPT1* has been proposed to participate in the establishment of a vascular neural stem cell niche within which quiescence and activation can be regulated^157^. In addition to pericytes which secrete *ANGPT1* to signal to endothelial cells, it is also expressed in neurons and has proneurogenic effects on subventricular zone stem/progenitor cells^158^. Notably, *ANGPT1* signals by binding to TIE2 receptors which in turn regulate mTOR activity (see mTOR below, this section)^158^.

Thus both dog and human GWASs of diverse behavioral traits are associated with neurogenesis and neurodevelopment. That resonates with the notion of a “p” factor. The psychiatric-disorder factors of internalization and externalization, once thought to be distinct and separate, are now known to be substantially correlated and therefore to share risk factors^159^. This led to the idea of a “p” factor that reflects risk for all psychopathology and two residual factors for internalization and externalization risk. That is supported by a large and rapidly growing body of evidence^160^. For instance, a recent human study generated models of psychopathology dimensions using data on 11 psychiatric disorders and 10 normative and non-normative personality traits in 2,796 Norwegian twins^161^. They interpreted three factors: a general risk factor for all psychopathology, a second specific for internalizing traits and a third specific for externalizing traits. Estimates of their heritability and 95% confidence intervals were 48% (41-54%), 35% (28-42%), and 37% (31-44%), respectively.

It is notable that canine body size loci are most strongly enriched for the IGF1 growth-signaling pathway^154^, which has major impacts on neurogenesis^155^. Another canine haplotype strongly associated with behavior and body size contains *HMGA2*, which has direct roles in transcriptional control of the cell cycle – including self-renewal *vs.* differentiation decisions in neural stem cells^7-10^. Notably, the pathways of IGF1 and HMGA2 converge to regulate mTOR, possibly the most important regulator of growth and sizes of cell, organs and whole animals^162^. [See discussion of lack of evidence of epistasis between these two loci in section B.13, but note this does not rule out their biochemical relationships referenced here.] Based on internal and external cellular signaling, such as those related to energy metabolism states (incl. IGF1), mTOR serves as a master orchestrator of growth (mTOR Complex 1), and survival and proliferation (mTOR Complex 2), and their integration^163^. Importantly here, mTOR has central roles in stemness and neurogenesis^164^. One established way Insulin/IGF1 and HMGA2 interact is that the Insulin/IGF1 pathway gene IGF2BP2 – which is one of the candidate genes in our previous^5^ and present canine behavioral GWASs – is a direct target of HMGA2. That signaling system is part of the LIN28/let-7 loop that serves as a master regulator of glucose metabolism^165^ and has been proposed to regulate mTOR by balancing the growth factor/PI3K and MYC pathways^166^. In addition to Insulin/IGF1, this signaling network includes inputs of WNT/Frizzled (which also integrate PI3K and MYC pathways through beta catenin), and other growth factors (which also integrate PI3K and MYC pathways via RAS) that work together to regulate mTOR activity. Thus, there is evidence this overall balancing of mTOR signaling can be mediated by interacting protein-microRNA pairs comprised of LIN28A/B and members of the let-7 microRNA family^166^, and that LIN28/let-7 are key regulators of stemness and neurogenesis^164,167,168^. While it is widely appreciated how mTOR is central to energy metabolism, body size, longevity and reproduction^169^, there is a rapidly growing body of evidence of its key roles in diverse aspects of brain function and disease^170-175^. This suggests a mechanistic explanation for the correlations between these traits.

Consistent with the suggestions in the preceding paragraph, the EMBL-EBI human GWAS Catalog lists LIN28A or LIN28B/LIN28B-AS1 as the mapped genes for the following traits: for LIN28A, height, appendicular lean mass, unipolar depression response to trauma exposure; and for LIN28B, height, multiple body fat traits, age at menarche and other puberty traits, trauma exposure, unipolar depression, depressive symptoms, schizophrenia, insomnia, neuroticism, multiple smoking behaviors, wellbeing, guilt measurement and cognitive decline in Alzheimer’s. That is strong support for broad inherited risks of psychopathology. It also indicates several variations involved in gene interactions with the environment – including traits related to diet/energy metabolism, tobacco use and trauma. Aside from such inherited risk, consideration of the LIN28/let-7 loop in the context of behavior seems particularly important for environmental effects more generally, including its role in the sensitive window of development associated with risk of psychopathology. One recent publication is consistent with this possibility, showing childhood traumatization and neuropsychiatric outcomes are associated with plasma levels of four microRNAs, including let-7g^176^. Circulating levels of let-7g are also abnormal in schizophrenia and a common polymorphism at its locus, in an interval spanning 30 genes, is associated with schizophrenia risk^177,178^. There is also human genetic and animal model evidence linking that LIN28/let-7 biology to pleiotropic effects of let-7 on energy metabolism, body size, longevity and reproduction^179,180^, and showing this is evolutionarily conserved from worms and flies to humans^181-184^.

It is not difficult to imagine how selection impacting mTOR biology could affect cancer risk. Associations between psychiatric traits, including autism, and cancer are well known and variations in the PI3K-Akt-mTOR axis are prominent^185^. There is a strong negative correlation between canine body size and longevity (*R*=−0.68, *P*=6.73E−10), but not between cancer mortality and longevity (*R*=−0.06, *P*=0.63)^186-188^. However, cancer mortality residuals corrected for the correlation between body size and cancer mortality showed a positive correlation of cancer mortality and longevity (*R*=0.24, *P*=0.05, with evidence this was not influenced by outliers)^186^. Several hypotheses arise from that and other observations in this work (e.g., those indicated by long-standing and newer theories of cancer risk^189-191^). The fact dog breeds carry diverse inherited variations associated with body size and cancer presents an ideal molecular-epidemiological and translational model for testing such hypotheses^59,92-94^.

**B.11. Dog coat color gene agouti (*ASIP*) has behavioral effects suggestive of domestication**

The subiculum, discussed above, is also specifically implicated by one of our strongest behavioral loci and its leading candidate gene, Agouti-signaling protein or *ASIP*. This locus was associated with nonsocial fear, familiar dog-directed aggression and separation anxiety in two cohorts, and other traits in single cohorts. *ASIP* was associated with energy in MacLean et al.’s gene-based association meta-analysis^40^; we also found this locus to be associated with energy in the Boyko cohort^2^ that was not included in MacLean et al. It should be noted that individual *ASIP*-allele assays are insufficient to provide complete genotype-phenotype associations and the SNP platforms used here do not have the necessary coverage to allow determination of that based on haplotypes^192^. We showed here (section A.3) that human *ASIP* brain expression is highest in the subiculum, followed by the central nucleus of the amygdala; both of those regions have key roles in fear and anxiety^193,194^. It was recently shown that inactivation of *ASIP* in a wild strain of mouse results in a domestication phenotype^195^. Tameness or domestication *Asip* effects were previously reported for mink, deer mouse and rat^196-199^. In 1942, Keeler wrote “Thus, it seems probable that the tame albino rat, at least the strain studied, was not domesticated by painstaking selection over long periods of time, but was modified in morphology principally by the introduction of three coat color genes, and in behavior particularly by the (non-agouti) black gene.”^200,201^. In 1968, he reported domestication traits exhibited by silver/black foxes – which were subsequently shown to be agouti null^202^ – in comparison to amber-colored WT red foxes (*Vulpes vulpes*)^203^. In 1959, Belyaev initiated the “Russian farm-fox experiment” that went on to select for tame foxes for over 50 generations^204^ (critically re-evaluated in Ref. ^205^). Notably, that founding population was descendant from a docile, silver/black fox population farmed in Prince Edward Island, Canada, since the late 1800’s^206^. The *ASIP* locus is under strong selection in diverse domesticated species (incl. dogs, sheep, goats and horses). However, despite evidence it also has major roles in energy metabolism^207^ and behavior^195-203^, there seems to be a longstanding consensus that *ASIP* has only been under selection for coat color and pattern^208^. However, our GWA findings are in agreement with Keeler, Hirose et al. and others who produced evidence that *Asip* deficiency has major domesticating effects. Our human and cattle datamining evidence is consistent with brain expression effects reported by Hirose et al. and suggest ASIP is not just a pigmentation signaling protein, but a brain neuropeptide. Given C57BL/6, the predominant mouse strain used in neuroscience, is null for *ASIP*, it is important to learn how that contributes to diverse biological phenomena.

*ASIP* was deleted from the genome early in the lineage of one of our closest primate relatives, the gibbon (est. 25MYA)^209^. It was proposed that the loss of *ASIP* could explain their remarkably small body mass among the apes^209^. Another uncommon trait of gibbons is their use of swinging from tree to tree as the predominant locomotor activity. This is dangerous, as evidenced by the high frequency of broken bones in gibbons^210,211^ – which leads us to speculate that loss of *ASIP* could also have been selected for behavioral effects, such as reduced fear.

**B.12. Implications of behavioral GWA loci containing multiple biologically-relevant candidate genes**

Our prior annotation of the chromosome 18 and X loci showed they both implicated multiple relevant genes with highly tissue-specific co-expression (*GNAT3* and *CD36*, and *ARHGAP36*, *IGSF1*, and *FIRRE,* respectively)^5^. Here we noted *RALY*, *EIF2S2* and *AHCY* at the chr24 locus are co-expressed in activated, but not quiescent, neural stem cells. Other loci were notable for having multiple genes with strong biological relevance. The following are examples from quasi-replicated loci: *MSRB3* and *HMGA2 on* chr10; *MITF* and *FOXP1* on chr20; *RALY*, *EIF2S2, ASIP* and *AHCY* on chr24, and *IGF2BP2* and *TRA2B* on chr34. These loci could contain variations affecting one gene or multiple variations affecting one or more genes. However, these examples raise the possibility that genetic variations at these loci could affect expression of multiple genes in the same genetic or biological pathway. Follow-on studies should consider that such loci could exhibit multi-gene dysregulation. For example, such effects could arise from mutation of a CTCF binding site or presence of a structural variant. However, even functional coding-variation, as is strongly implicated for the behavior and body size *IGSF1* risk allele^5,23^, could result in feedback-regulation affecting multiple genes at this enigmatic locus that also contains 15 CTCF binding sites and the lncRNA *FIRRE*, which is a central regulator of non-homologous chromosomal contacts^212,213^.

**B.13. Epistasis analysis of quasi-replicated loci**

Even the simplest epistatic interaction analysis is very computationally demanding when the full genome is included. In addition, the analysis of epistasis generates very large amounts of output that complicate their interpretation. Very conservative thresholds to declare significance are not enough to limit the amount of data generated since the sensitivity of the statistical tools and the inherent likelihood of detecting interactive effects is relatively large. To mitigate those challenges and anticipating follow-on studies of genetic testing for diagnosis and risk prediction, we designed the epistatic analysis to focus on our prioritized GWAS main-effect loci. Although this design sacrifices detecting potential markers outside of the GWAS relevant loci, we consider it a reasonable tradeoff that made the interpretation of the data more intuitive. Another challenge was the variation in datasets: differences in array platforms, allele call algorithms, quality control steps, and sample-related differences such as sample size and breed representation. This complicated direct comparisons and generalized use of specific threshold values.

We encountered additional challenges in the epistasis analysis with the Hayward dataset. It was designed to evaluate specific pathology risk and may have had higher representation of variations in specific breeds. It also contained a far larger sample size in comparison to the other two datasets. These factors are likely to have caused detection issues related to overpowered analysis. Although the Hayward cohort was genotyped with the Illumina Canine HD array with added SNPs, the number of SNPs available was smaller than expected in comparison to the Vaysse cohort which used the same array without the added SNPs. We believe a difference in the allele calling algorithm and quality control could be responsible for this issue since we observed many markers missing in the Hayward data were located in previously defined CNV regions^214,215^. At early stages of our analysis, we explored the use of a reduced dataset that would match the size of the other cohorts. We abandoned the idea after we observed in the GWAS that the reduced dataset modified the hit detection in a way we could not predict. Thus that approach could not be easily incorporated for the interpretation across all cohorts. It also violated our idea of providing direct comparisons of independent analysis of three independent cohorts with minimal data manipulation. Other factors that could affect the epistasis analysis are population structure and the effects of alleles that are fixed in many breeds. One possibly-mitigating factor is that the loci tested are enriched for interbreed admixture directly selected by humans for morphology and possibly behavior (i.e., vs. representing phylogenetic breed relationships)^216^. With the caveats above, we included that analysis here as an exploratory or preliminary first point. These should only be considered as nominated interactions requiring further support.

The largest-effect body size variations in this work were at the chr15 *IGF1* locus and a complex region of chr10 spanning *MSRB3* and *HMGA2*. We noted above that IGF1 and HMGA2 converge in mTOR biochemistry. Based on our threshold for epistasis, those two loci did not interact. However, they both interacted with the chr24 locus spanning *RALY*, *EIF2S2, ASIP* and *AHCY* , and the chrX locus containing a protein-coding variant in *IGSF1* that is presumed to be the functional variant for large body size^12^ and reduced risk of several problem behaviors^5^. The chrX/*IGSF1* locus also interacted with the chr13/*ANGPT1* locus. Several of these candidate interactions can be tested. Chr15 *IGF1* and chrX/*IGSF1* suggest IGF1 signaling and regulation of organismal growth by the hypothalamus and pituitary gland. As we mentioned gene co-expression at several loci that could be associated with epigenetic regulation in the previous section and this pathway is implicated in longevity^217^, we note that two candidate interacting-loci contain methionine metabolism genes, *MSRB3* and *AHCY*. Chr13/*ANGPT1* and chr18/*CD36* suggest neurovascular physiology. Several candidates hint at other possible epistatic effects such as neurogenesis (*IGF1*, *MITF* and *ANGPT1*) and stemness (*HMGA2*, *FOXP1* and *ANGPT1*). Alternatively, variation in expression levels or tissue specificity of candidate gene alleles could also suggest testable hypotheses.

**B.14. Conclusions**

We propose behavioral adaptive evolution preferentially targets a specific genetic network. It is associated with neurogenesis and neurodevelopment, and suggests a molecular basis for pleiotropy between behavior and height^82,85,112,218^, BMI/energy metabolism^218^, reproduction^180,219^ and longevity^220^. The core evolutionary selection involved here (under the top-most drivers, survival and reproduction) appears to involve the balance between energy metabolism and growth/development. This is consistent with Witting’s theory^221^ and draws attention to the effects of domestication of animals and human self-domestication. That and the possible impact on cancer risk are addressed above (section B.10). Our analyses suggest we were able to detect these effects with breed stereotypes of behavior and moderately-powered genesets because of the large effect sizes of dog variations under strong selection and under greatly relaxed negative selection. This would not be possible in similarly powered studies of humans, which do not have similar genetic architecture for the traits under study here. However, our cross-species findings for behavioral traits show strong genetic conservation.

We showed many transcription factors appear to have central roles in the super-network implicated by canine behavioral mapping, including subfamily members of the Fox (incl. FOXO, FOXA and FOXD) and Hox (incl. POU, LHX, MLX, ALX, CUX and PAX) families, as well as NR2C2/F2, SMAD2/4 and MEF2A. In section B.10 above we cite the evidence for nominating mTOR and its regulation by the ancient LIN28/let-7 loop as a central hub of the pleiotropic behavioral biology discussed here. That is supported with respects to behavior by common LIN28A/B variations associated with inherited behavioral and psychiatric traits. The evidence for that includes gene-environment interactions, such as exposure to trauma. That suggests the possibility this biochemistry could be affected by stress during the critical neurodevelopmental time window that is associated with increased psychiatric disease risk (i.e., in general, with or without increased genetic susceptibility).

Multiple theories have been developed to explain how neurogenesis could be targeted in the evolutionary adaptation of behavior (without linking that to the “p” factor)^222^. Unlike what we propose here, those theories tend to be rather complex and hinge on fine details of subtypes of neuroplasticity and memory functions, exact developmental sites and time-points of adult neurogenesis, and specific behaviors. Many have proposed that human evolutionary adaptation is associated with psychiatric disease risks (without linking that to the p factor)^223^. While research articles have increasingly highlighted the implication of neurogenesis and neurodevelopment pathways in human psychiatric genetics^5,40,80-86,142,143^, the same authors^142^ can omit the terms neurogenesis and neurodevelopment in a review titled “Psychiatric genetics and the structure of psychopathology”^224^ (the terms are also absent from the list of theories for the “p” factor in Discussion of ref. ^161^). We believe a general genetic factor of psychopathology (correlated with personality and intelligence^225,226^) is likely to involve neurodevelopment from fetal through the full life course. However, specific disorders can have earlier or later developmental effects or disease onset, such as neurodevelopmental disorders and autism being enriched for embryonic neurodevelopment and having an early onset. Humans have strongly selected not only for canine working traits and morphology, but also for companionship and personality. This suggests brain function and its relationships to body-wide physiology can be genetically-dissected much more readily in dogs compared to humans.

**C. Expanded methods**

**C.1. Direct phasing of SNP based genotypes**

Direct phasing was performed as previously described^227^. Briefly, the selected marker and sorted by breed and their carrier status. Only homozygous individuals are further grouped by their allele (A/B). For each group, markers upstream and downstream are sequentially evaluated and only kept if they had an allele frequency over 0.95. This provides homozygous block segmentation that can be visualized across breeds.

**C.2. Epistasis analysis**

Epistatic interactions were only assessed among the GWAS hits under the significance threshold for their respective dataset. All epistatic analysis were performed in PLINK v.1.07. Significance was declared for this analysis also by using a Bonferroni adjusted thresholds calculated for each data set by only considering the number of valid tests performed. Since the goal of this study was to detect relevant interactions among markers that would be suggestive of specific physiological mechanisms that controls behavior, we based the interpretation of epistatic interactions on the number of significant interactions among markers across all traits. To visualize epistatic data side to side across datasets we developed circular plots using the CIRCOS v.0.69 application^228^.

**D. Expanded references**

199.	Price, E.O. *Animal domestication and behavior*, (Cabi, 2002).

200.	Keeler, C.E. The association of the black (non-agouti) gene with behavior: In the Norway Rat. *Journal of Heredity* **33**, 371-384 (1942).

201.	Keeler, C.E. & King, H.D. Multiple effects of coat color genes in the Norway rat, with special reference to temperament and domestication. *Journal of Comparative Psychology* **34**, 241 (1942).

202.	Vage, D.I. *et al.* A non-epistatic interaction of agouti and extension in the fox, Vulpes vulpes. *Nat Genet* **15**, 311-5 (1997).

203.	Keeler, C.E. *Behavior synthesis through pigment gene pleiotropy*, (1968).

204.	Kukekova, A.V., Temnykh, S.V., Johnson, J.L., Trut, L.N. & Acland, G.M. Genetics of behavior in the silver fox. *Mamm Genome* **23**, 164-77 (2012).

205.	Lord, K.A., Larson, G., Coppinger, R.P. & Karlsson, E.K. The History of Farm Foxes Undermines the Animal Domestication Syndrome. *Trends Ecol Evol* **35**, 125-136 (2020).

206.	Petersen, M. *The fur traders and fur bearing animals*, (Hammond Press, 1914).

**Supplementary figures**

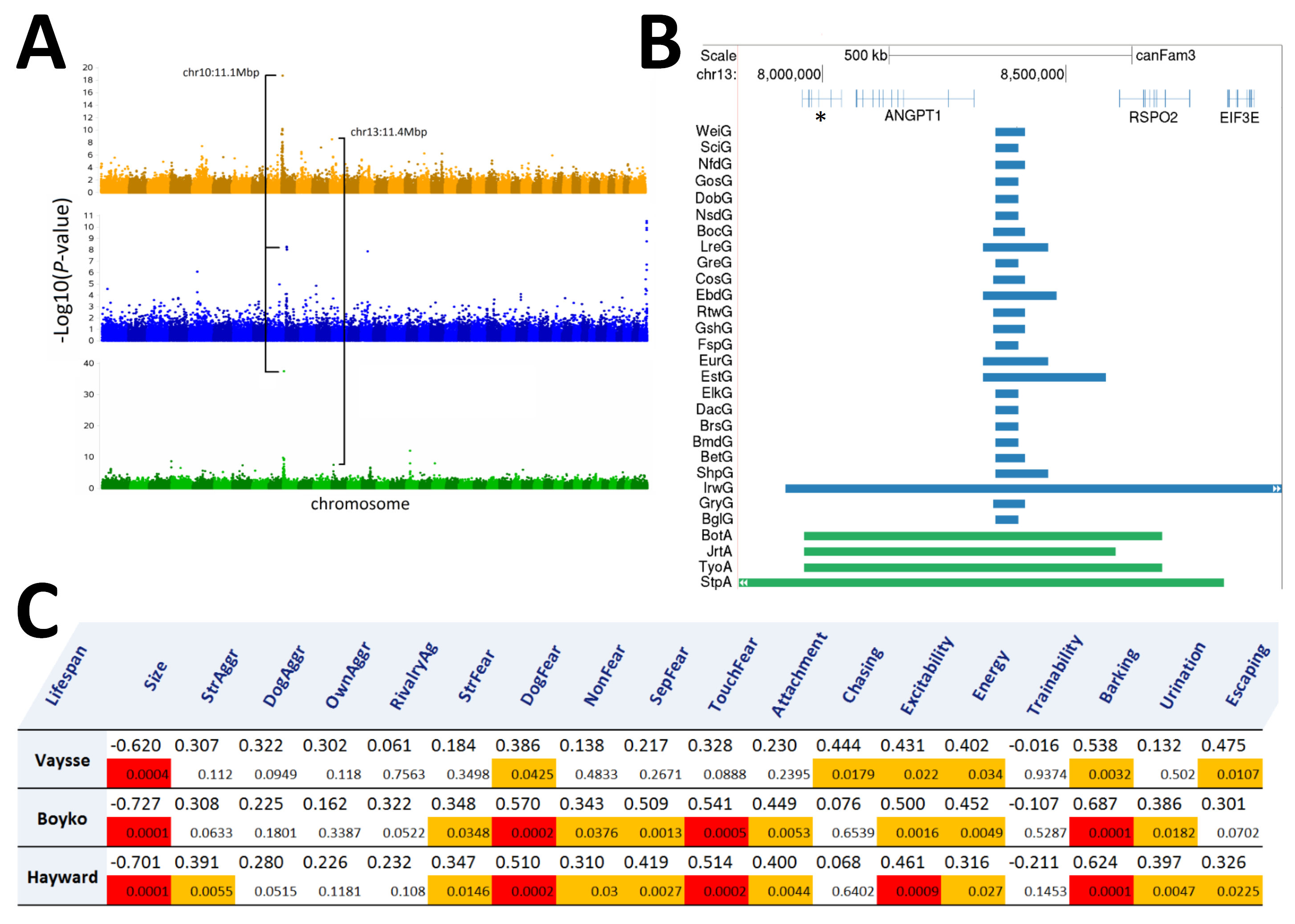

**Figure S1. Expected lifespan GWASs on three independent cohorts.** A) Manhattan plots of Lifespan displaying regions where GWAS hits were replicated using a Bonferroni corrected threshold that was independently calculated for each dataset. Vaysse (top), Boyko (center) and Hayward (bottom). B) Direct phasing and mapping of Vaysse SNPs by breed on the fixed position Chr13:8,391,652 (CanFam3.1) * Refers to a uncharacterized predicted protein ENSCAFG00000000687. C) Pearson’s Correlation coefficients and P-values for Lifespan phenotypes vs. Size and problematic behavior traits. P-values highlighted in orange are ≤0.05, while those in red are Bonferroni corrected.

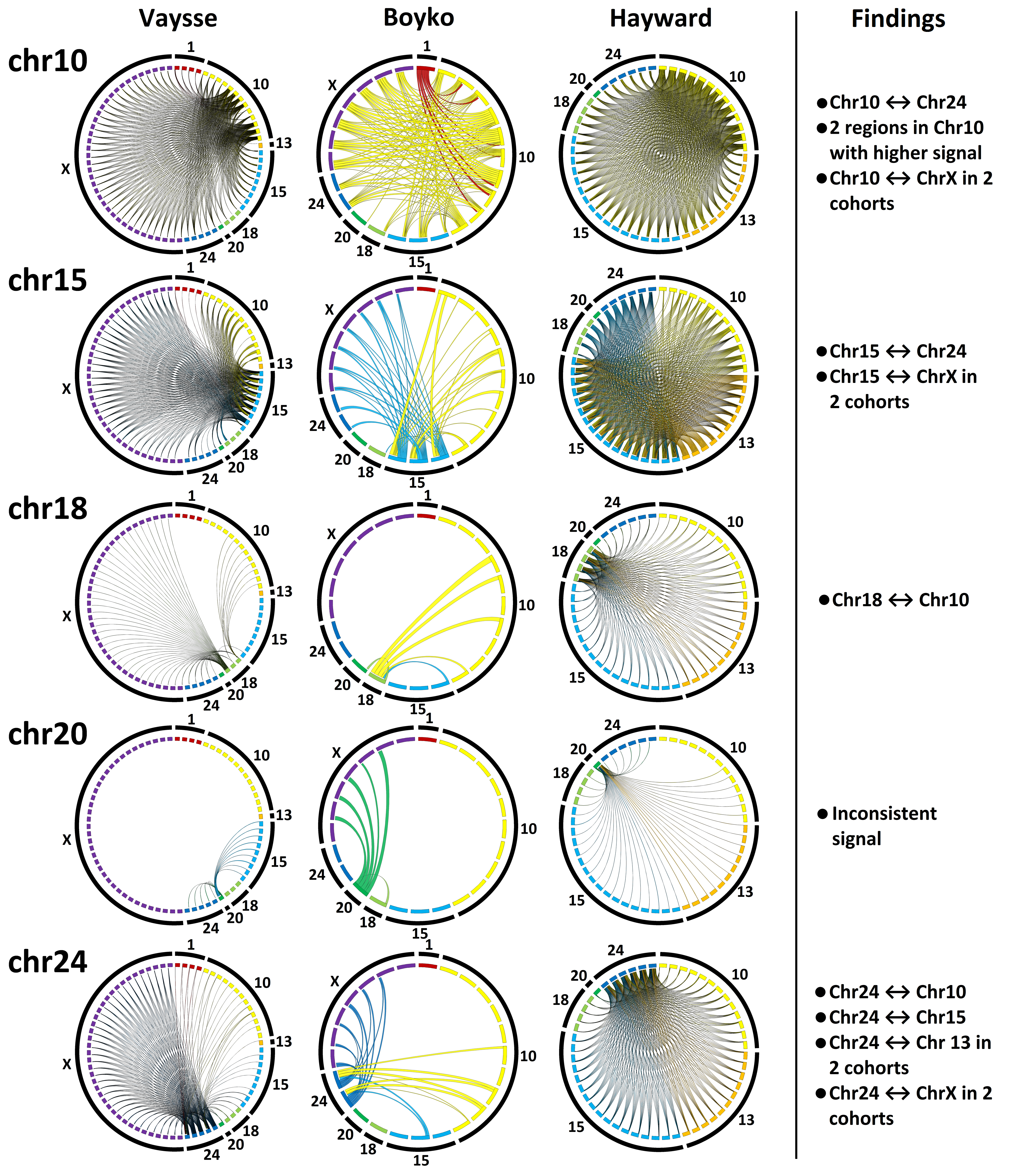

**Figure S2. Epistasis.** CIRCOS plots of epistatic interactions among replicated GWAS hits of problematic behavior in dog traits across three independent cohorts. Columns correspond to Vaysse, Boyko and Hayward. Only significant hits using a Bonferroni corrected threshold were included in the analysis. Epistatic interactions among adjacent markers in the same locus is not displayed since it is directly related to LD. Ribbon color indicates the origin of the comparison and its thickness indicates the number of times a significant interaction was detected among all traits. Only the epistatic interactions significant to a Bonferroni adjusted threshold are displayed. Markers are displayed in genomic order clockwise.
